# Decomposing variability in protein levels from noisy expression, genome duplication and partitioning errors during cell-divisions

**DOI:** 10.1101/026559

**Authors:** M. Soltani, C. A. Vargas-Garcia, D. Antunes, A. Singh

## Abstract

Inside individual cells, expression of genes is inherently stochastic and manifests as cell-to-cell variability or noise in protein copy numbers. Since proteins half-lives can be comparable to the cell-cycle length, randomness in cell-division times generates additional intercellular variability in protein levels. Moreover, as many mRNA/protein species are expressed at low-copy numbers, errors incurred in partitioning of molecules between the mother and daughter cells are significant. We derive analytical formulas for the total noise in protein levels for a general class of cell-division time and partitioning error distributions. Using a novel hybrid approach the total noise is decomposed into components arising from i) stochastic expression; ii) partitioning errors at the time of cell-division and iii) random cell-division events. These formulas reveal that random cell-division times not only generate additional extrinsic noise but also critically affect the mean protein copy numbers and intrinsic noise components. Counter intuitively, in some parameter regimes noise in protein levels can decrease as cell-division times become more stochastic. Computations are extended to consider genome duplication, where the gene dosage is increased by two-fold at a random point in the cell-cycle. We systematically investigate how the timing of genome duplication influences different protein noise components. Intriguingly, results show that noise contribution from stochastic expression is minimized at an optimal genome duplication time. Our theoretical results motivate new experimental methods for decomposing protein noise levels from single-cell expression data. Characterizing the contributions of individual noise mechanisms will lead to precise estimates of gene expression parameters and techniques for altering stochasticity to change phenotype of individual cells.

## 1 Introduction

The level of a protein can deviate considerably from cell-to-cell, in spite of the fact that cells are genetically-identical and are in the same extracellular environment [1–3]. This intercellular variation or noise in protein counts has been implicated in diverse processes such as corrupting functioning of gene networks [4–6], driving probabilistic cell-fate decisions [7–12], buffering cell populations from hostile changes in the environment [13–16], and causing clonal cells to respond differently to the same stimulus [17–19]. An important source of noise driving random fluctuations in protein levels is stochastic gene expression due to the inherent probabilistic nature of biochemical processes [20–23]. Recent experimental studies have uncovered additional noise sources that affect protein copy numbers. For example, the time take to complete cell-cycle (i.e., time between two successive cell-division events) has been observed to be stochastic across organisms [24–32]. Given that many proteins/mRNAs are present inside cells at low-copy numbers, errors incurred in partitioning of molecules between the mother and daughter cells are significant [33–35]. Finally, the time at which a particular gene of interest is duplicated can also vary between cells [36,37]. We investigate how such noise sources in the cell-cycle process combine with stochastic gene expression to generate intercellular variability in protein copy numbers (Fig.1).

**Figure 1.**
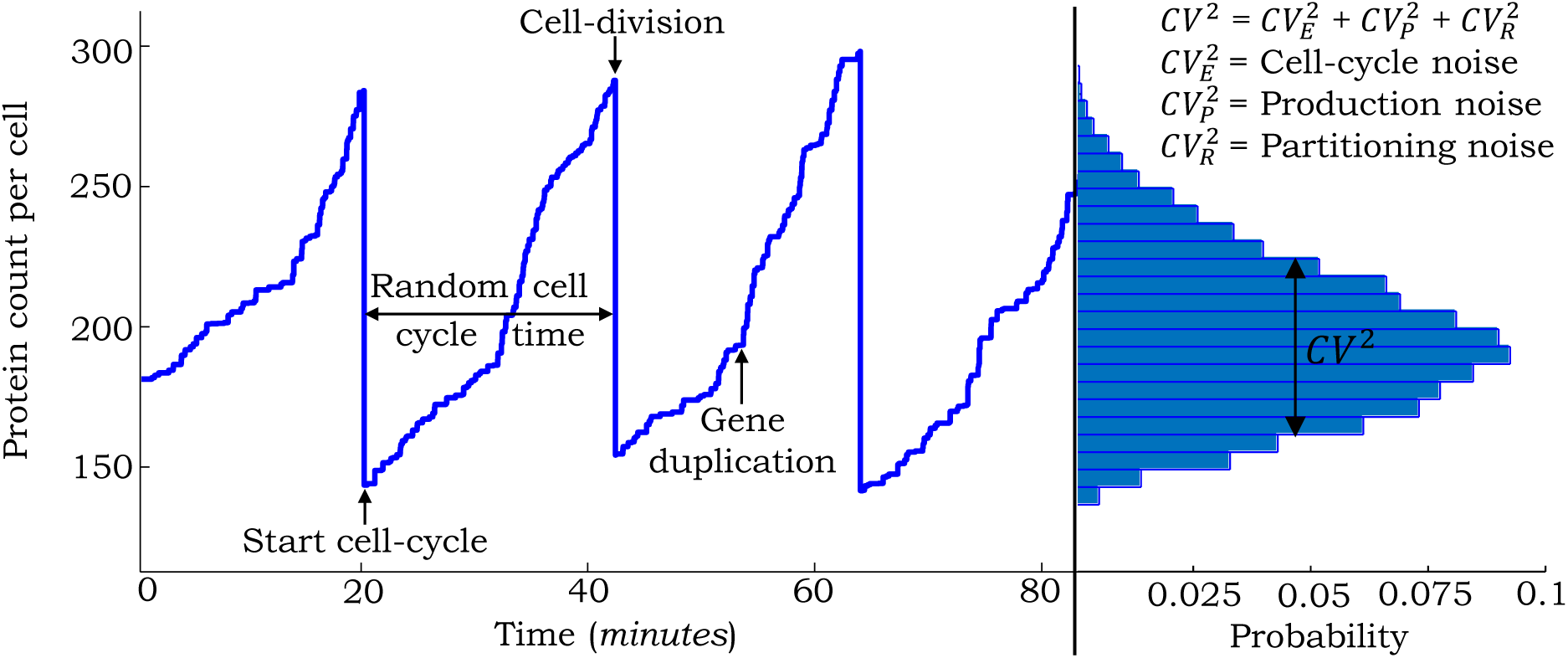
Sample trajectory of the protein level in a single cell with different sources of noise. Stochastically expressed proteins accumulate within the cell at a certain rate. At a random point in the cell-cycle, gene-duplication results in an increase in production rate. Stochastic cell-division events lead to random partitioning of protein molecules between the mother and daughter cells with each cell receiving, on average, half the number of proteins in the mother cell just before division. The steady-state protein copy number distribution obtained from a large number of trajectories is shown on the right. The total noise in the protein level, as measured by the Coefficient of Variation (*CV*) squared can be broken into contributions from individual noise mechanisms.

Prior studies that quantify the effects of cell-division on the protein noise level have been restricted to specific cases. For example, noise computations have been done in stochastic gene expression models, where cell-divisions occur at deterministic time intervals [33,38,39]. Recently, we have analyzed a deterministic model of gene expression with random cell-division events [40]. Building up on this work, we formulate a mathematical model that couples stochastic expression of a stable protein with random cell-division events that follow an arbitrary probability distribution function. Moreover, at the time of cell-division, proteins are randomly partitioned between the mother and daughter cells based on a general framework that allows the partitioning errors to be higher or lower than as predicted by binomial partitioning. For this class of models, we derive an exact analytical formula for the protein noise level as quantified by the steady-state Coefficient of Variation (*CV*) squared. This formula is further decomposed into individual components representing contributions from different noise sources. A systematic investigation of this formula leads to novel insights, such as identification of regimes where increasing randomness in the timing of cell-division events decreases the protein noise level.

Next, we extend the above model to include genome duplication events that increase the gene’s transcription rate by two-fold (corresponding to doubling of gene dosage) prior to cell-division [36,41]. To our knowledge, this is the first study integrating randomness in the genome duplication process with stochastic gene expression. An exact formula for the protein noise level is derived for this extended model and used to investigate how the timing of duplication affects different noise components. Counter intuitively, results show that doubling of the transcription rate within the cell-cycle can lead to smaller fluctuations in protein levels as compared to a constant transcription rate through out the cell-cycle. Finally, we discuss how formulas obtained in this study can be used to infer parameters and characterize the gene expression process from single-cell studies.

## 2 Coupling gene expression to cell-division

We consider the standard model of stochastic gene expression [42,43], where mRNAs are transcribed at exponentially distributed time intervals from a constitutive gene with rate *k_x_*. For the time being, we exclude genome duplication and the transcription rate is fixed throughout the cell-cycle. Assuming short-lived mRNAs, each transcription event results in a burst of proteins [43–45]. The corresponding jump in protein levels is shown as

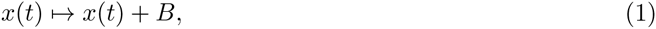

where *x*(*t*) is the protein population count in the mother cell at time *t*, *B* is a random burst size drawn from a positively-valued distribution and represents the number of protein molecules synthesized in a single-mRNA lifetime. Motivated by observations in *E. coli* and mammalian cells, where many proteins have half-lives considerably longer than the cell-doubling time, we assume a stable protein with no active degradation [46–48]. Thus, proteins accumulate within the cell till the time of cell-division, at which point they are randomly partitioned between the mother and daughter cells.

Let cell-division events occur at times *t_s_*, *s* ∈ {1, 2,…}. The cell-cycle time

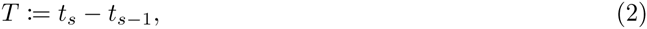

follows an arbitrary positively-valued probability distribution with the following mean and coefficient of variation (*CV*) squared

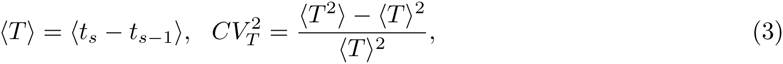

where 〈.〉 denotes expected value through out this paper. The random change in *x*(*t*) during cell-division is given by

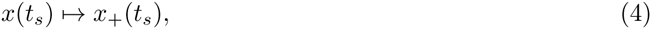

Where *x*(*t_s_*) and *x*_+_(*t_s_*) denote the protein levels in the mother cell just before and after division, respectively. Conditioned on *x*(*t_s_*), *x*_+_(*t_s_*) is assumed to have the following statistics

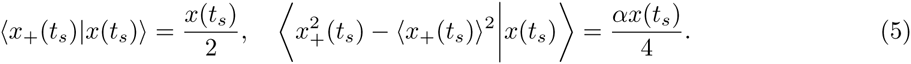

The first equation implies symmetric division, i.e., on average the mother cell inherits half the number protein molecules just before division. The second equation in (5) describes the variance of 〈*x*_+_(*t_s_*)〉 and quantifies the error in partitioning of molecules through the non-negative parameter α. For example, *α* = 0 represents deterministic partitioning where *x*_+_(*t_s_*) = *x*(*t_s_*)/2 with probability equal to one. A more realistic model for partitioning is each molecule having an equal probability of being in the mother or daughter cell [49–51]. This result in a binomial distribution for *x*_+_(*t_s_*)

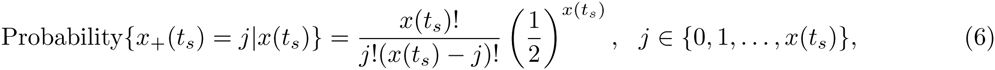

and corresponds to *α* = 1 in (5). Interestingly, recent studies have shown that partitioning of proteins that form clusters or multimers can result in *α* > 1 in (5), i.e., partitioning errors are much higher than as predicted by the binomial distribution [33,39]. In contrast, if molecules push each other to opposite poles of the cell, then the partitioning errors will be smaller than as predicted by (6) and *α* < 1.

The model with all the different noise mechanisms (stochastic expression; random cell-division events and partitioning errors) is illustrated in Fig. 2A and referred to as the full model. We also introduce two additional hybrid models [52,53], where protein production and partitioning are considered in their deterministic limit (Fig. 2B-C). Note that unlike the full model, where *x*(*t*) takes non-negative integer values, *x*(*t*) is continuous in the hybrid models. We will use these hybrid models for decomposing the protein noise level obtained from the full model into individual components representing contributions from different noise sources. However, before computing the noise, we first determine the average number of proteins as a function of the cell-cycle time distribution.

**Figure 2.**
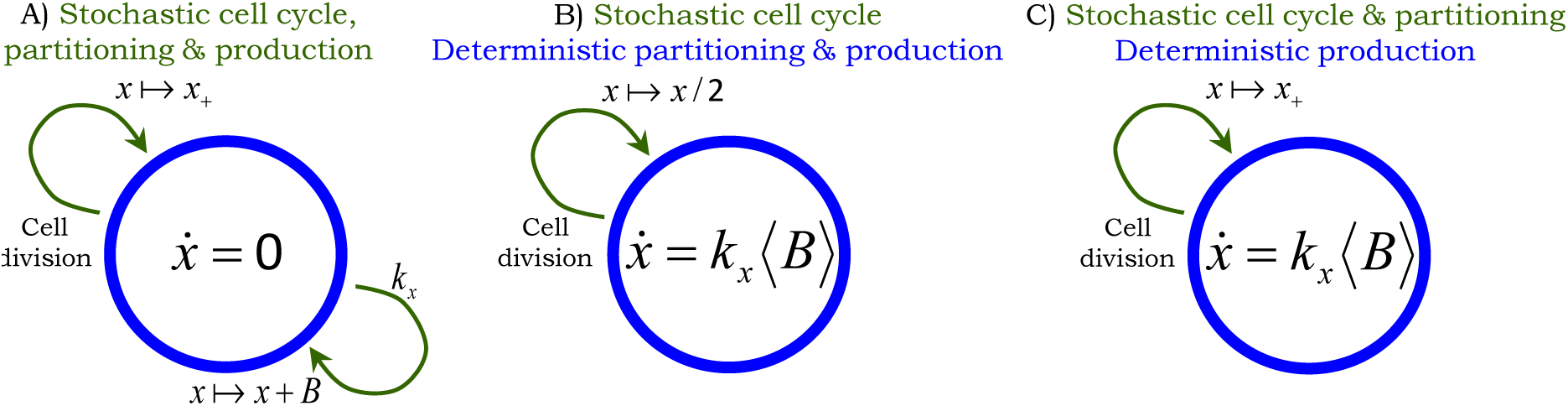
Stochastic models of gene expression with cell-division. Arrows denote stochastic events that change the protein level by discrete jumps as shown in (1) and (4). The differential equation within the circle represents the time evolution of *x*(*t*) in between events. **A)** Model with all the different sources of noise: proteins are expressed in stochastic bursts, cell-division occurs at random times, and molecules are partitioned between the mother and daughter cells based on (5). The trivial dynamics *x* = 0 signifies that the protein level is constant in-between stochastic events. **B)** Hybrid model where randomness in cell-division events is the only source of noise. Protein production is modeled deterministically through a differential equation and partitioning errors are absent, i.e., *α* = 0 in (5). **C)** Hybrid model where noise comes from both cell-division events and partitioning errors. Protein production is considered deterministically as in Fig. 2B. Since *x*(*t*) is continuous here, *x*_+_(*t_s_*) has a positively-valued continuous distribution with same mean and variance as in (5)

## 3 Computing the average number of protein molecules

To quantify the steady-state mean protein level we consider the full model illustrated in Fig. 2A. It turns out that all the models shown in Fig. 2 are identical in terms of finding 〈*x*(*t*)〉 and in principle any one of them could have been used. To obtain differential equations describing the time evolution of 〈*x*(*t*)〉 we model the cell-cycle time through a phase-type distribution, which can be represented by a continuous-time Markov chain. Phase-type distributions are dense in the class of positively-valued continuous distributions, i.e., one can always construct a sequence of phase-type distributions that converges point wise to a given distribution of interest [54]. We use this denseness property as a practical tool for modeling the cell-cycle time.

### 3.1 Cell-cycle time as a phase-type distribution

We consider a class of phase-type distribution that consists of a mixture of Erlang distributions. Recall that an Erlang distribution of order *i* is the distribution of the sum of *i* independent and identical exponential random variables. The cell-cycle time is assumed to have an Erlang distribution of order *i* with probability *p_i_*, *i* = {1,…, *n*} and can be represented by a continuous-time Markov chain with states *G_ij_*, *j* = {1,…, *i*}, *i* = {1,…, *n*} (Fig. 3). Let Bernoulli random variables *g_ij_* = 1 if the system in the next infinitesimal resides in state *G_ij_* and 0 otherwise. The probability of transition *G_ij_* → *G_i_*_(_*_j_* _+ 1)_, time interval [*t*, *t* + *dt*) is given by *kg_ij_dt*, implying that the time spent in each state *G_ij_* is exponentially distributed with mean 1/*k*. To summarize, at the start of cell-cycle, a state *G_i_*_1_, *i* = {1,…, *n*} is chosen with probability *p_i_* and cell-division occurs after transitioning through *i* exponentially distributed steps. Based on this formulation, the probability of a cell-division event occurring in the next time interval [*t*, *t* + *dt*) is given by 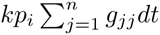, and whenever the event occurs, the protein level changes as per (4). Finally, the mean and the coefficient of variation squared of the cell-cycle time is obtained as

**Figure 3.**
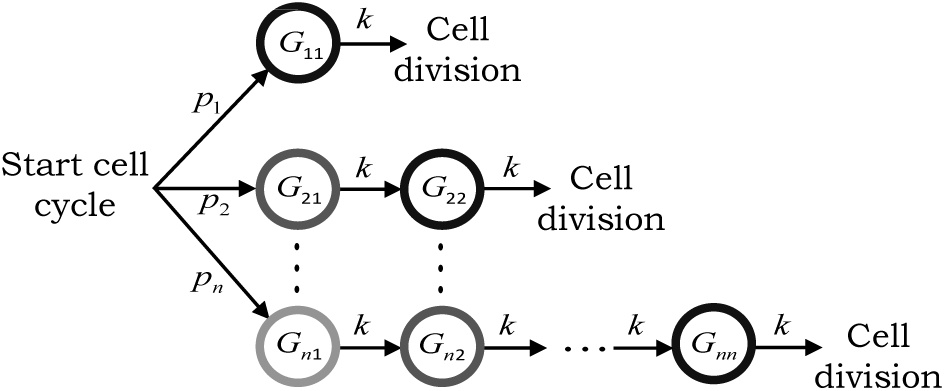
A continuous-time Markov chain model for the cell-cycle time. The cell-cycle time is assumed to follow a mixture of Erlang distributions. At the start of cell-cycle, a state *G_i_*_1_, *i* = {1,…, *n*} is chosen with probability *p_i_*,. The cell-cycle transitions through states *G_i_*_j_, *j* = {1,…, *i*} residing for an exponentially distributed time with mean 1/*k* in each state. Cell-division occurs after exit from *G_ii_* and the above process is repeated.

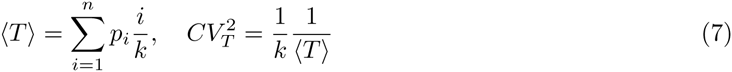

in terms of the Markov chain parameters. Our goal is to obtain 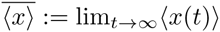 as a function of 〈*T*〉 and 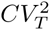.

### 3.2 Time evolution of the mean protein level

Time evolution of the statistical moments of *x*(*t*) can be obtained from the Kolmogorov forward equations corresponding to the full model in Fig. 2A combined with the cell-division process described in Fig. 3. We refer the reader to [52, 55, 56] for an introduction to moment dynamics for stochastic and hybrid systems. Analysis in Appendix A shows

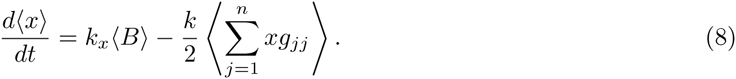

Note that the time-derivative of the mean protein level (first-order moment) is unclosed, in the sense that, it depends on the second-order moment 〈*xg_ij_*〉. Typically, approximate closure methods are used to solve moments in such cases [52,56–61]. However, the fact that *g_ij_* is binary can be exploited to automatically close moment dynamics. In particular, since *g_ij_* ∈ {0,1}

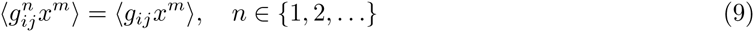

for any non-negative integer *m*. Moreover, as only a single state *g_ij_* can be 1 at any time

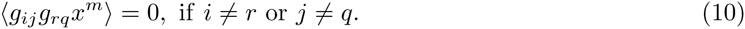

Using (9) and (10), the time evolution of 〈*xg_ij_*〉 is obtained as

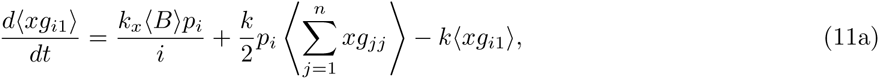

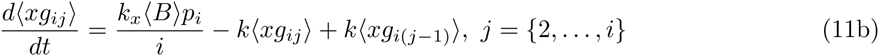

and only depends on 〈*xg_ij_*〉 (see Appendix A). Thus, (8) and (11) constitute a closed system of linear differential equations from which moments can be computed exactly.

To obtain an analytical formula for the average number of proteins, we start by performing a steadystate analysis of (8) that yields

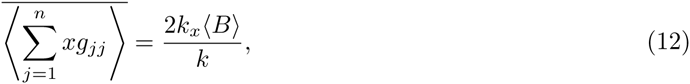

where 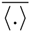 denotes the expected value in the limit *t* → ∞. Using (12), 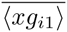 is determined from (11a), and then all moments 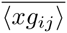 are obtained recursively by performing a steady-state analysis of (11b) for *j* = {2,…, *i*}. This analysis results in

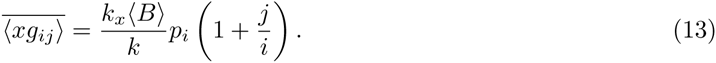

Using (7), (13) and the fact that 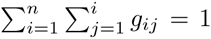 we obtain the following expression for the mean protein level

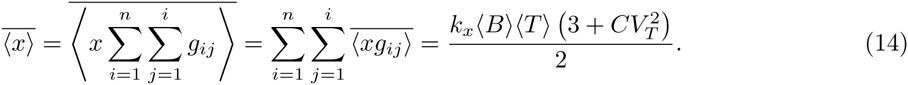

It is important to point that (14) holds irrespective of the complexity, i.e., the number of sates *G_ij_* used in the phase-type distribution to approximate the cell-cycle time distribution. As expected, 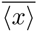 increases linearly with the average cell-cycle time duration 〈*T*〉 with longer cell-cycles resulting in more accumulation of proteins. Consistent with previous findings, (14) shows that the mean protein level is also affected by the randomness in the cell-cycle times 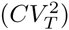 [40,62]. For example, 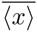 reduces by 25% as *T* changes from being exponentially distributed 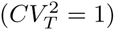 to periodic 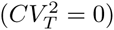 for fixed 〈*T*〉 fixed. Next, we determine the noise in protein copy numbers, as quantified by the coefficient of variation squared.

## 4 Computing the protein noise level

Recall that the full model introduced in Fig. 2A has three distinct noise mechanisms. Our strategy for computing the protein noise level is to first analyze the model with a single noise source, and then consider models with two and three sources. As shown below, this approach provides a systematic dissection of the protein noise level into components representing contributions from different mechanisms.

### 4.1 Contribution from randomness in cell-cycle times

We begin with the model shown in Fig. 2B, where noise comes from a single source - random cell-division events. For this model, the time evolution of the second-order moment of the protein copy number is obtained as

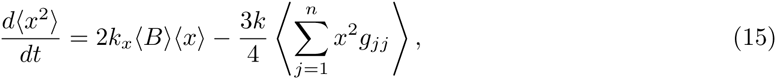

and depends on third-order moments 〈*x*^2^*g_jj_*〉 (see Appendix B). Using the approach introduced earlier for obtaining the mean protein level, we close moment equations by writing the time evolution of moments 〈*x*^2^*g_ij_*〉. Using (9) and (10)

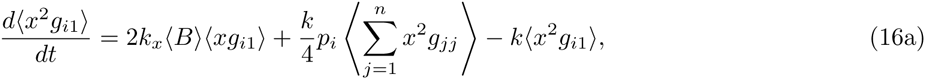

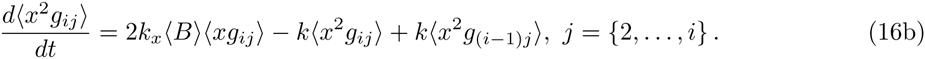

Note that the moment dynamics for 〈*x*〉 and 〈*xg_ij_*〉 obtained in the previous section (equations (8) and (11)) are identical for all the models in Fig. 2, irrespective of whether the noise mechanism is modeled deterministically or stochastically. Equations (8), (11), (15) and (16) represent a closed set of linear differential equations and their steady-state analysis yields

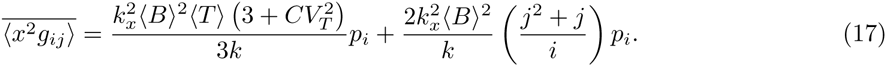

From (96)

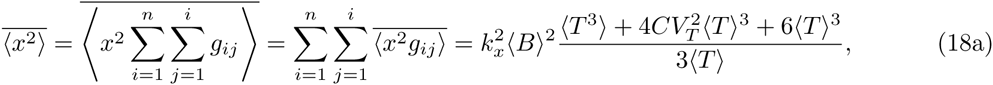

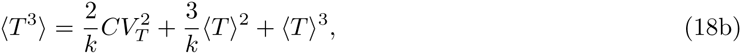

where 〈*T*^3^〉 is the third-order moment of the cell-cycle time. Using (18) and the mean protein count quantified in (14), we obtain the following coefficient of variation squared

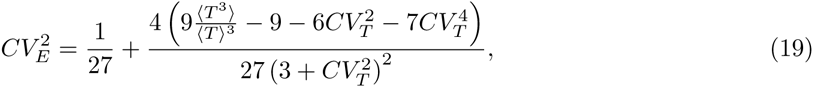

which represents the noise contribution from random cell-division events. Since cell-division is a global event that affects expression of all genes, this noise contribution can also be referred to as *extrinsic noise* [49,63–66]. In reality, there would be other sources of extrinsic noise, such as, fluctuations in the gene-expression machinery that we have ignored in this analysis.

Note that 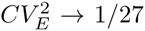 as *T* approaches a delta distribution, i.e., cell divisions occur at fixed time intervals. We discuss simplifications of (19) in various limits. For example, if the time taken to complete cell-cycle is lognormally distributed, then

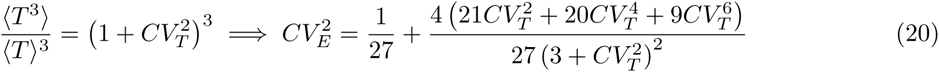

and extrinsic noise monotonically increases with 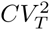. If fluctuations in *T* around 〈*T*〉 are small, then using Taylor series

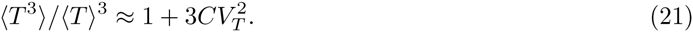

Substituting (21) in (19) and ignoring 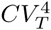 and higher order terms yields

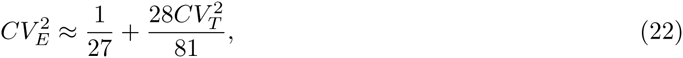

where the first term is the extrinsic noise for 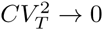 and the second term is the additional noise due to random cell-division events.

### 4.2 Contribution from partitioning errors

Next, we consider the model illustrated in Fig. 2C with both random cell-division events and partitioning of protein between the mother and daughter cells. Thus, the protein noise level here represents the contribution from both these sources. Analysis in Appendix C shows that the time evolution of 〈*x*^2^〉 and 〈*x*^2^*g_ij_*〉 are given by

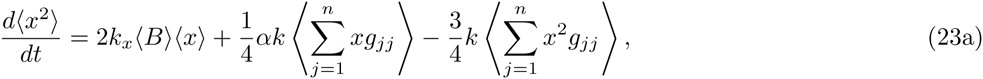

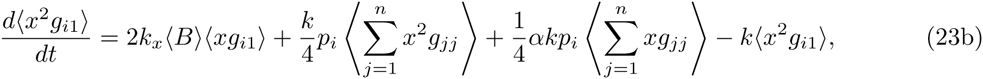

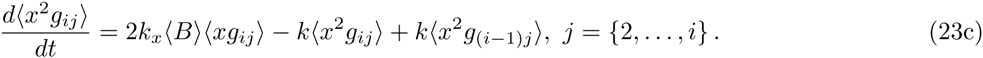

Note that (23a)-(23b) are slightly different from their counterparts obtained in the previous section (equations (15) and (16a)) with additional terms that depend on *α*, where *α* quantifies the degree of partitioning error as defined in (5). As expected, (23) reduces to (15)-(16) when *α* = 0 (i.e., deterministic partitioning). Computing 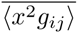 by performing a steady-state analysis of (23) and using a similar approach as in (18) we obtain

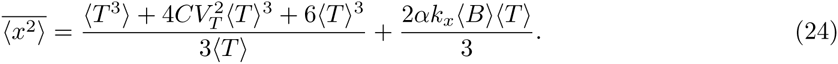

Finding *CV*^2^ of the protein level and subtracting the extrinsic noise found in (19) yields

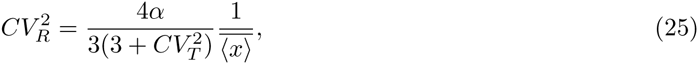

where 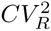 represents the contribution of partitioning errors to the protein noise level. Intriguingly, while 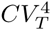 increases with *α*, it decrease with 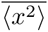. Thus, as cell-division times become more random for a fixed 〈*T*〉 and 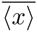, the noise contribution from partitioning errors decrease.

### 4.3 Contribution from stochastic expression

Finally, we consider the full model in Fig. 2A with all the three different noise sources. For this model, moment dynamics is obtained as (see Appendix D)

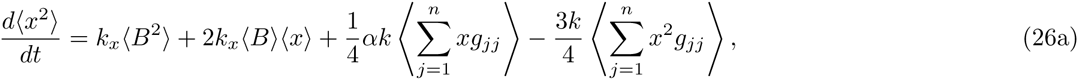

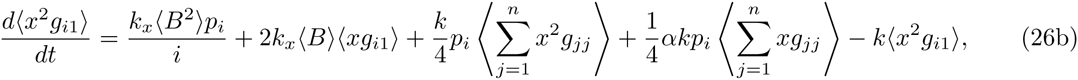

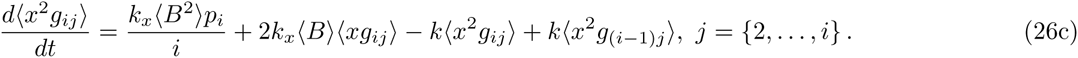

Compared to (23), (26) has additional terms of the form *k_x_*〈*B*^2^〉, where 〈*B*^2^〉 is the second-order moment of the protein burst size in (1). Performing an identical analysis as before we obtain

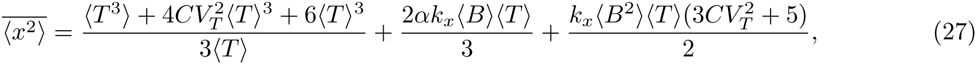

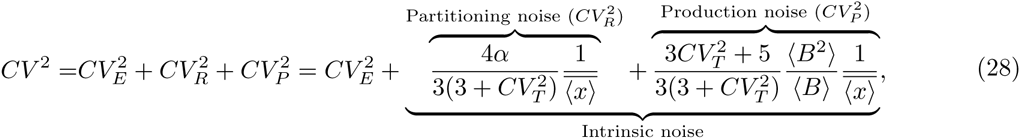

that can be decomposed into three terms. The first is the extrinsic noise 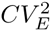 representing the contribution from random cell-division events and given by (19). The second term 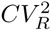 is the contribution from partitioning errors determined in the previous section (partitioning noise), and the final term 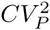 is the additional noise representing the contribution from stochastic expression (production noise). We refer to the sum of the contributions from partitioning errors and stochastic expression as *intrinsic noise.* These intrinsic and extrinsic noise components are generally obtained experimentally using the dual-color assay that measures the correlation in the expression of two identical copies of the gene [49].

Interestingly, for a fixed mean protein level 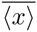, 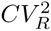 has opposite effects on 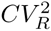 and 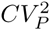 While 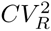 monotonically decreases with increasing 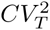, 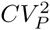 increases with 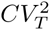. It turns out that in certain cases these effects can cancel each other out. For example, when *B* = 1 with probability one, i.e., proteins are synthesized one at a time at exponentially distributed time intervals and *α* = 1 (binomial partitioning)

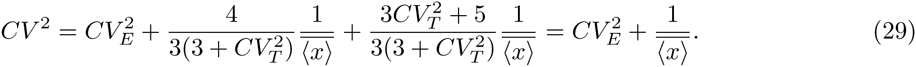

In this limit the intrinsic noise is always 1/Mean irrespective of the cell-cycle time distribution *T* [33]. Note that the average number of proteins itself depends on *T* as shown in (14). Another important limit is 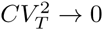, in which case (28) reduces to

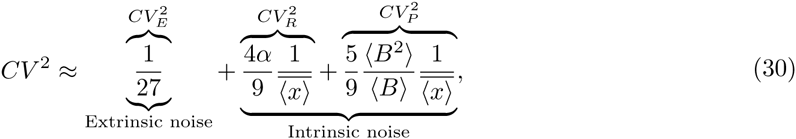

and is similar to the result obtained in [38] for deterministic cell-division times and binomial partitioning.

Fig. 4 shows how different protein noise components change as a function of the mean protein level as the gene’s transcription rate *k_x_* is modulated. The extrinsic noise is primarily determined by the distribution of the cell-cycle time and is completely independent of the mean. In contrast, both 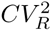 and 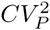; scale inversely with the mean, albeit with different scaling factors (Fig. 4). This observation is particularly important since many single-cell studies in *E. coli,* yeast and mammalian cells have found the protein noise levels to scale inversely with the mean across different genes [67–70]. Based on this scaling it is often assumed that the observed cell-to-cell variability in protein copy numbers is a result of stochastic expression. However, as our results show, noise generated thorough partitioning errors is also consistent with these experimental observations and it may be impossible to distinguish between these two noise mechanisms based on protein *CV*^2^ versus mean plots unless *α* is known.

**Figure 4.**
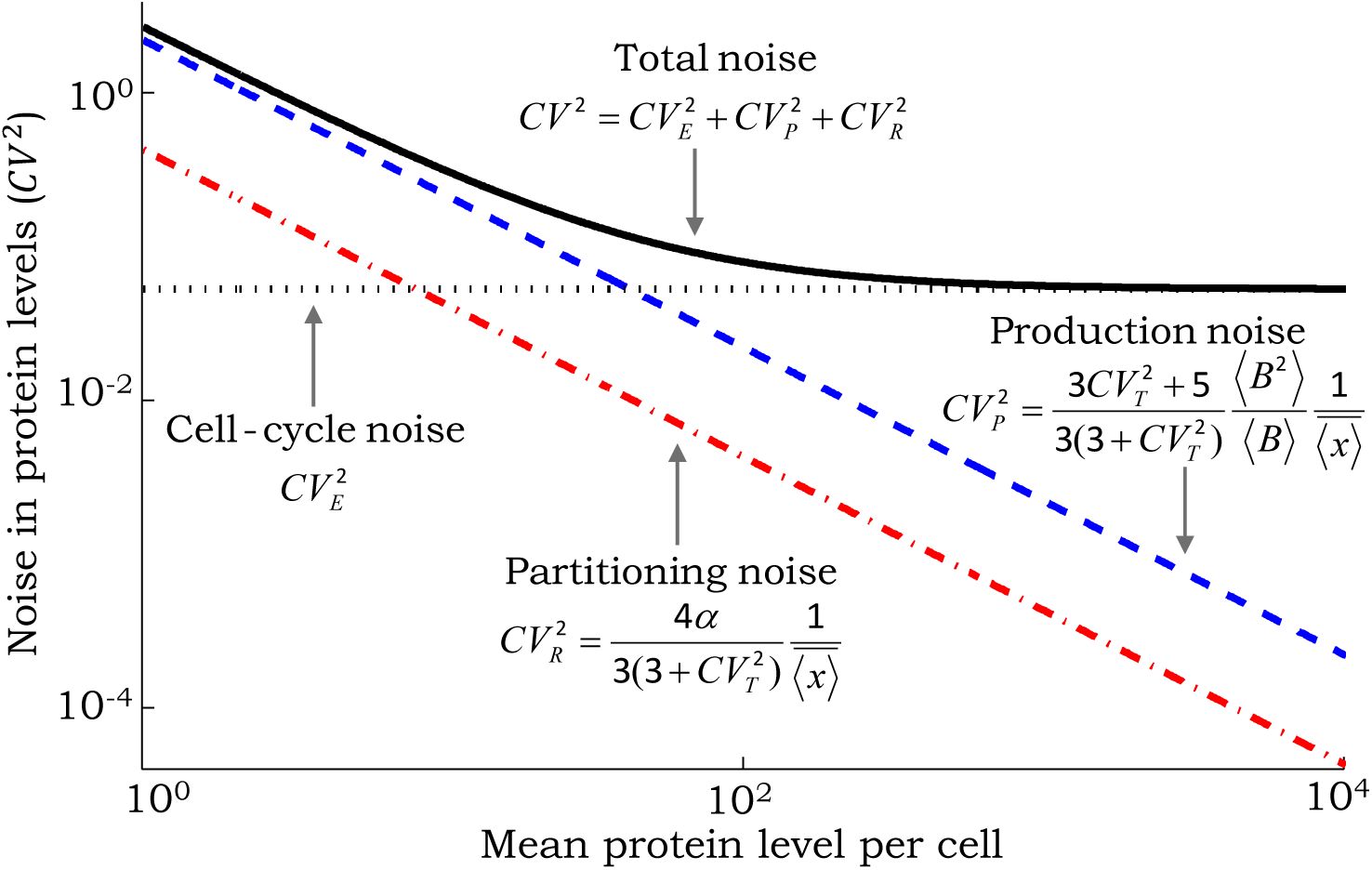
Scaling of noise as a function of the mean protein level for different mechanisms. The contribution of random cell-division events to the noise in protein copy numbers (extrinsic noise) is invariant of the mean. In contrast, contributions from partitioning errors at the time of cell-division (partitioning noise) and stochastic expression (production noise) scale inversely with the mean. The scaling factors are shown as a function of the protein random burst size *B*, noise in cell-cycle time 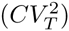 and magnitude of partitioning errors quantified by *α* (see (5)). With increasing mean level the total noise first decreases and then reaches a baseline that corresponds to extrinsic noise. For this plot, *B* is assumed to be geometrically-distributed with mean 〈*B*〉 = 1.5, 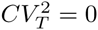 and *α* = 1 (i.e., binomial partitioning).which yields the following total protein noise level

## 5 Quantifying the effects of gene-duplication on protein noise

The full model introduced in Fig. 2 assumes that the transcription rate (i.e., the protein burst arrival rate) is constant throughout the cell-cycle. This model is now extended to incorporate gene duplication during cell cycle, which is assumed to create a two-fold change in the burst arrival rate (Fig. 5). As a result of this, accumulation of proteins will be bilinear as illustrated in Fig. 1. We divide the cellcycle time *T* into two intervals: time from the start of cell-cycle to gene-duplication (*T*_1_), and time from gene-duplication to cell-division (*T*_2_). *T* and *T*_2_ are independent random variables that follow arbitrary distributions modeled through phase-type processes (see Fig. S2 in the Supplementary Information). The mean cell-cycle duration and its noise can be expressed as

**Figure 5.**
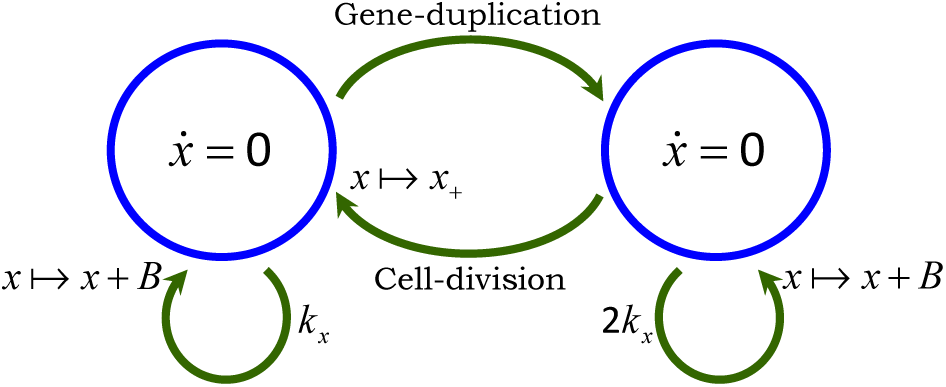
Model illustrating stochastic expression together with random geneduplication and cell-division events. At the start of cell-cycle, protein production occurs in stochastic bursts with rate *k_x_*. Genome duplication occurs at a random point *T*_1_ within the cellcycle and increases the burst arrival rate to 2*k_x_*. Cell-division occurs after time *T*_2_ from genome duplication, at which point the burst arrival rate reverts back to and proteins are randomly partitioned between cells based on (4).

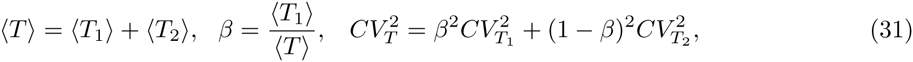

Where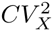denotes the coefficient of variation squared of the random variable *X*. An important variable in this formulation is *β*, which represents the average time of gene-duplication normalized by the mean cell-cycle time. Thus, *β* values close to 0 (1) imply that the gene is duplicated early (late) in the cell-cycle process. Moreover, the noise in the gene-duplication time is controlled via 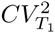.

We refer the reader to Appendix E for a detailed analysis of the model in Fig. 5 and only present the main results on the protein mean and noise levels. The steady-state mean protein count is given by

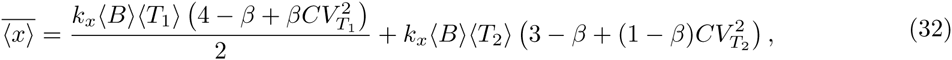

and decreases with *β*, i.e., a gene that duplicates early has on average, more number of proteins. When *β* = 1, then the transcription rate *k_x_* is throughout the cell-cycle and we recover the mean protein level obtained in (14). Similarly, when *β* = 0 the transcription rate is 2*k_x_* and we obtain twice the amount as in (14). As per our earlier observation, more randomness in the timing of genome duplication and cell-division (i.e., higher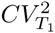and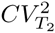values) increases 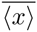.

Our analysis shows that the total protein noise level can be decomposed into three components

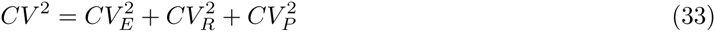

where 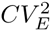 is the extrinsic noise from random genome-duplication and cell-division events. Given its complexity, we refer the reader to equation (100) in Appendix E2 for an exact formula for 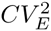. Moreover, the intrinsic noise, which represents the sum of contributions from partitioning errors 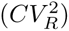 and stochastic expression 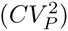 is obtained as

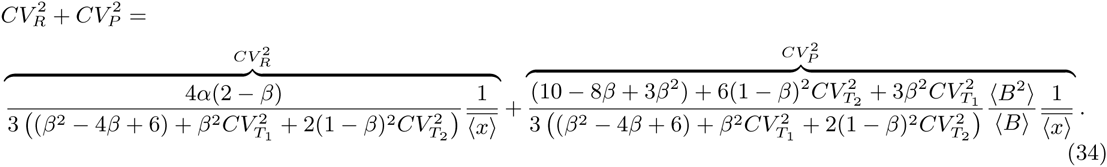

Note that for *β* = 0 and 1, we recover the intrinsic noise level in (28) from (34). Interestingly, for *B* = 1 with probability 1 and *α* = 1, the intrinsic noise is always 1/Mean irrespective of the values chosen for 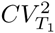, 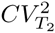and *β*. For high precision in the timing of cell-cycle events 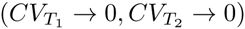

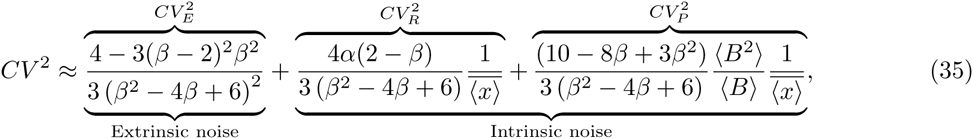

where mean protein level is given by

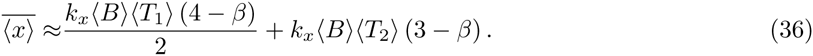

We investigate how different noise components in (35) vary with 0 as the mean protein level is held fixed by changing *k_x_*. Fig. 6 shows that 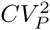 follows a U-shaped profile with the optima occurring at 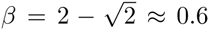 and the corresponding minimum value being ≈ 5% lower that its value at *β* = 0. An implication of this result is that if stochastic expression is the dominant noise source, then geneduplication can result in slightly lower protein noise levels. In contrast to 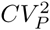, 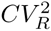 has a maxima at 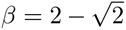 which is ≈ 6% higher than its value at *β* = 0 (Fig. 6). Analysis in Appendix E5 reveals that 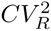 and 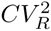 follow the same qualitative shapes as in Fig. 6 for non-zero 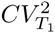 and 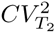. Interestingly, when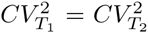, the maximum and minimum values of 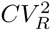 and 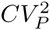 always occur at 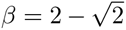 albeit with different optimal values than Fig. 6 (see Fig. S3 in the Supplementary Information). For example, if 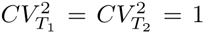 (i.e., exponentially distributed *T*_1_ and *T*_2_), then the maximum value of 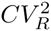 is 20% higher and the minimum value of 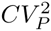 is 10% lower than their respective value for *β* = 0. Given that the effect of changing *β* on 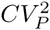 and 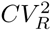 is small and antagonistic, the overall affect of genome duplication on intrinsic noise may be minimal and hard to detect experimentally.

**Figure 6.**
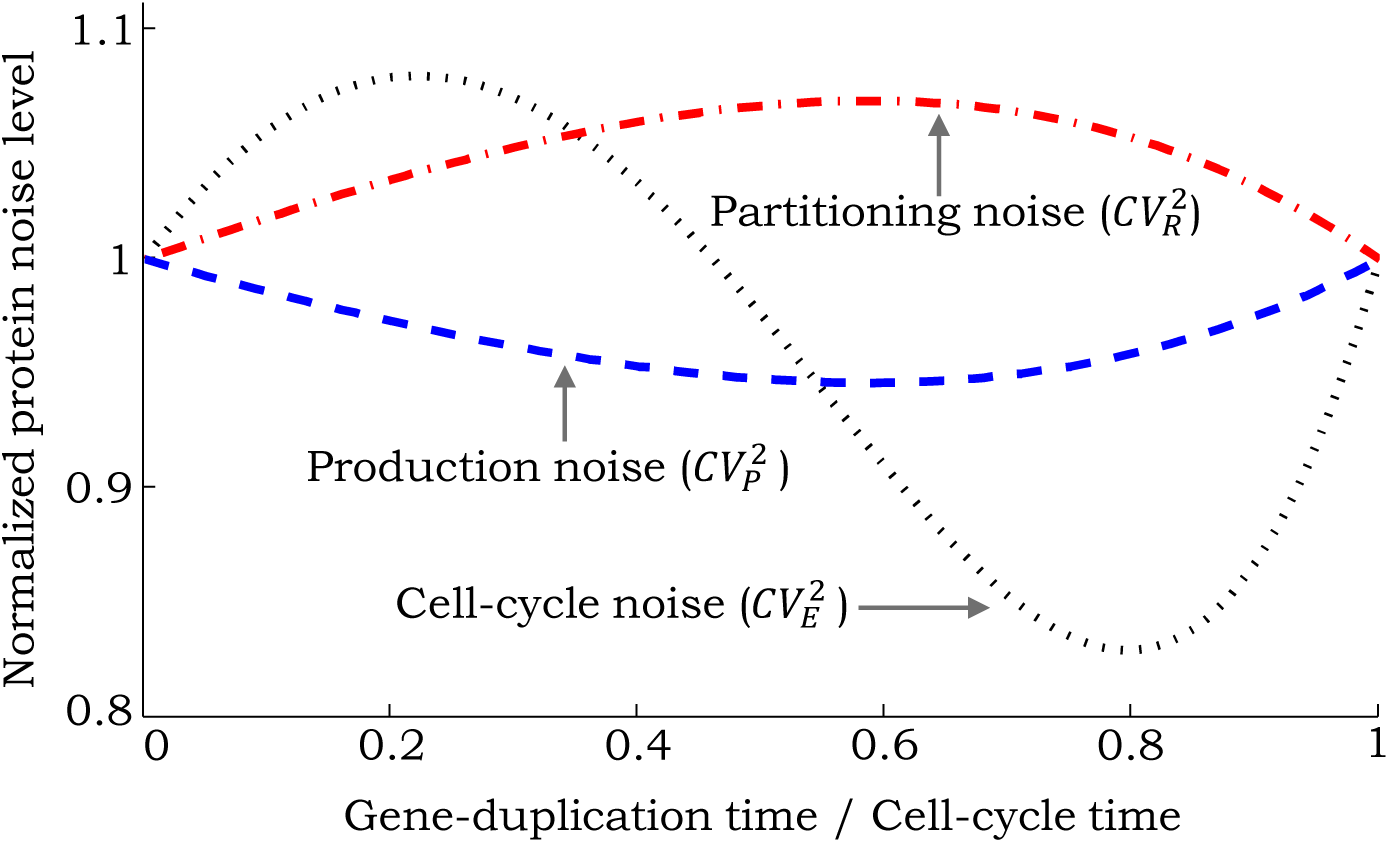
Contributions from different noise sources as a function of the timing of genome duplication for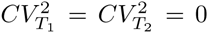. Different noise components in (35) are plotted as a function of *β*, which represents the fraction of time within the cell-cycle at which gene-duplication occurs. The mean protein level is held constant by simultaneously changing the transcription rate *k_x_*. Noise levels are normalized by their respective value at *β* = 0. The noise contribution from partitioning errors is maximized at *β* ≈ 0.6. In contrast, the contribution from stochastic expression is minimum at *β* ≈ 0.6. The extrinsic noise contribution from random gene-duplication and cell-division events is maximum at *β* ≈ 0.2 and minimum at *β* ≈ 0.8.

## 6 Discussion

We have investigated a model of protein expression in bursts coupled to discrete gene-duplication and cell-division events. The novelty of our modeling framework lies in describing the size of protein bursts, *T*_1_ (time between cell birth and gene duplication), *T*_2_ (time between gene duplication and cell division) and partitioning of molecules during cell division through arbitrary distributions. Exact formulas connecting the protein mean and noise levels to these underlying distributions were derived. Furthermore, the protein noise level, as measured by the coefficient of variation squared, was decomposed into three components representing contributions from gene-duplication/cell-division events, stochastic expression and random partitioning. While the first component is independent of the mean protein level, the other two components are inversely proportional to it. Key insights obtained are as follows:

- The mean protein level is affected by both the first and second-order moments of *T*_1_ and *T*_2_. In particular, randomness in these times (for a fixed mean) increases the average protein count.
- Random gene-duplication/cell-division events create an extrinsic noise term which is completely determined by moments of *T*_1_ and *T*_2_ up to order three.
- The noise contribution from partitioning errors decreases with increasing randomness in *T*_1_ and *T*_2_. Thus, if 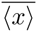 is sufficiently small and *α* is large compared to *B* in (34), increasing noise in the timing of cell-cycle events decreases the total noise level.
- Genome duplication has counter intuitive effects on the protein noise level (Fig. 6). For example, if stochastic expression is the dominant source of noise, then doubling of transcription due to duplication results in lower noise as compared to constant transcription throughout the cell-cycle.1.1e
- For a non-bursty protein production process (*B* = 1) and binomial partitioning (*α* = 1), the net noise from stochastic expression and partitioning is always 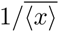, the noise level predicted by a Poisson distribution.

We discuss our results on gene duplication in further detail and how noise formulas derived here can be used for estimating model parameters from single-cell expression data.

### 6.1 Affect of gene duplication on noise level

In this first-of-its-kind study, we have investigated how discrete two-fold changes in the transcription rate due do gene duplication affect the intercellular variability in protein levels. Not surprisingly, the timing of genome duplication has a strong effect on the mean protein level 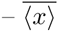 changes by two-fold depending on whether the gene duplicates early (*β* = 0) or late (*β* = 1) in the cell-cycle. In contrast, the effect of *β* on noise is quite small. As *β* is varied keeping 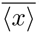 fixed, noise components deviate by ≈ 10% from their values at *β* = 0 (Fig. 6). Recall that these results are for a stable protein, whose intracellular copy number accumulate in a bilinear fashion. A natural question to ask is how would these results change for an unstable protein?

Consider an unstable protein with half-life considerably shorter than the cell-cycle duration. This rapid turnover ensures that the protein level equilibrates instantaneously after cell-division and geneduplication events. Let γ*_x_* denote the protein decay rate. Then, the mean protein level before and after genome duplication is 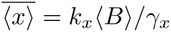 and 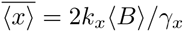, respectively. Note that in the limit of large *γ_x_* there is no noise contribution form partitioning errors since errors incurred at the time of cell division would be instantaneously corrected. The extrinsic noise, which can be interpreted as the protein noise level for deterministic protein production and decay is obtained as (see Appendix F)

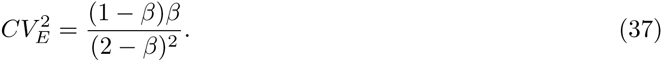

When *β* = 0 or 1, the transcription rate and the protein level are constant within the cell cycle and 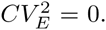. Moreover, 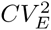 is maximized at *β* = 2/3 with a value of 1/12. Thus, in contrast to a stable protein, extrinsic noise in an unstable protein is strongly dependent on the timing of gene duplication. Next, consider the intrinsic noise component. Analysis in Appendix F shows that the noise contribution from random protein production and decay is

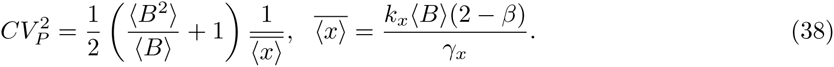

While the mean protein level is strongly dependent on *β*, the intrinsic noise Fano factor 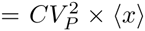 is independent of it. Thus, similar to what was observed for a stable protein, the intrinsic noise in an unstable protein is invariant of *β* for a fixed 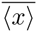. Overall, these results suggest that studies quantifying intrinsic noise in gene expression models, or using intrinsic noise to estimate model parameters (see below) can ignore the effects of gene duplication. Finally, note that the mean and noise levels obtained for an unstable protein are independent of the cell-cycle time *T*.

### 6.2 Parameter inference from single-cell data

Simple models of bursty expression and decay predict the distribution of protein levels to be negative binomial (or gamma distributed in the continuous framework) [71,72]. These distributions are characterized by two parameter – the burst arrival rate *k_x_* and the average burst size 〈*B*〉, which can be estimated from measured protein mean and noise levels. This method has been used for estimating and (B) across different genes in *E. coli* [47,73]. Our detailed model that takes into account partitioning errors predicts (ignoring gene duplication effects)

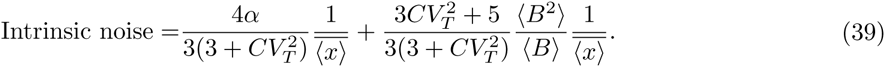

Using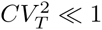 and a geometrically distributed *B* [50,74–76], (39) reduces to

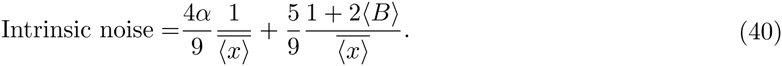

Given measurements of intrinsic noise and the mean protein level, 〈*B*〉 can be estimated from (40) assuming *α* = 1 (i.e., binomial partitioning). Once 〈*B*〉 is known, is obtained from the mean protein level given by (14). Since for many genes 〈*B*〉 ≈ 0.5 − 5 [47], the contribution of the first term in (40) is significant, and ignoring it could lead to overestimation of 〈*B*〉. Overestimation would be even more severe if *α* happen to be much higher than 1, as would be the case for proteins that form aggregates or multimers [33]. One approach to estimate both 〈*B*〉 and *α* is to measure intrinsic noise changes in response to perturbing 〈*B*〉 by, for example, changing the mRNA translation rate through mutations in the ribosomal-binding sites (RBS). Consider a hypothetical scenario where the Fano Factor (intrinsic noise times the mean level) is 6. Let mutations in the RBS reduces 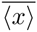 by 50%, implying a 50% reduction in (B). If the Fano factor changes from 6 to 4 due to this mutation, then 〈*B*〉 = 3.6 and 〈*α*〉 = 3.25.

Our recent work has shown that higher-order statistics of protein levels (i.e., skewness and kurtosis) or transient changes in protein noise levels in response to blocking transcription provide additional information for discriminating between noise mechanisms [77,78]. Up till now these studies have ignored noise sources in the cell-cycle process. It remains to be seen if such methods can be used for separating the noise contributions of partitioning errors and stochastic expression to reliably estimate 〈*B*〉 and *α*.

### 6.3 Integrating cell size and promoter switching

An important limitation of our modeling approach is that it does not take into account the size of growing cells. Recent experimental studies have provided important insights into the regulatory mechanisms controlling cell size [79–81]. More specifically, studies in *E. coli* and yeast argue for an “adder” model, where cell-cycle timing is controlled so as to add a constant volume between cell birth and division [82–84]. Assuming exponential growth, this implies that the time taken to complete cell-cycle is negatively correlated with cell size at birth. In addition, cell size also affects gene expression – in mammalian cells transcription rates linearly increase with the cell size [85]. Thus, as cells become bigger they also produce more mRNAs to ensure gene product concentrations remains more or less constant. An important direction of future work would to explicitly include cell size with size-dependent expression and timing of cell division determined by the adder model. This formulation will for the first time, allow simultaneous investigation of stochasticity in cell size, protein molecular count and concentration.

Our study ignores genetic promoter switching between active and inactive states, which has been shown to be a major source of noise in the expression of genes across organisms [86–95]. Taking into account promote switching is particularly important for genome duplication studies, where doubling the number of gene copies could lead to more efficient averaging of promoter fluctuations. Another direction of future work will be to incorporate this addition noise source into the modeling framework and investigate its contribution as a function of gene-duplication timing.

## Appendix

### A Mean of protein in the presence of cell-cycle variations

Based on standard stochastic formulation of chemical kinetics [96,97], the model introduced in Figure 2A coupled with phase-type distribution introduced in Figure 3 contains the following stochastic events

| Event | Reset | Propensity |
| --- | --- | --- |
| Protein production | $x(t) \mapsto x(t) + u$ | $k_x p_u''$ |
| Phase-type Evolution | $g_{ij}(t) \mapsto g_{ij}(t) - 1,$<br>$g_{i(j+1)}(t) \mapsto g_{i(j+1)}(t) + 1$ | $kg_{ij},$<br>$i \in \{2, \dots, n\},$<br>$j \in \{1, \dots, i-1\}$ |
| Cell-division | $x(t_s) \mapsto x_+(t_s),$<br>$g_{jj}(t_s) \mapsto 0,$<br>$g_{i1}(t_s) \mapsto g_{i1}(t_s) + 1$ | $kp_i \sum_{j=1}^n g_{jj},$<br>$i \in \{1, \dots, n\}$ |

Note that *x*_+_(*t_s_*) is protein level after division, characteristics of *x*_+_(*t_s_*) is related to protein level before division as shown in equation (5) of the main text. Whenever an event occurs, protein level and states of phase-type distribution change based on the stoichiometries shown in the second column of the table. The third column of table shows event propensity function *f*(*x*, *g_ij_*), which determines how often reactions occur, i.e., the probability that an event occurs in the next infinitesimal time interval (*t*, *t*+*dt*] is *f*(*x*, *g_ij_*)*dt*. Protein production is a stochastic event which happens in bursts, each burst generates *B* molecules where *B* is a general random variable with distribution

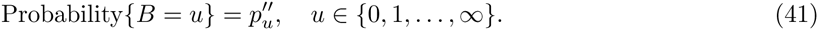

The probability of having a burst in the time interval (*t*, *t* + *dt*] is 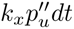. Events related to time evolution of phase-type distribution happen with a constant rate *k*. Cell-division changes both the level of protein and states of phase-type. This event contains start of new cell-cycle, hence whenever this event occurs, the last state of phase-type distribution resets to zero, and a new cell-cycle which is sum of *i* exponentials starts with probability *p_i_*; protein count level also resets to *x*_+_(*t_s_*). The probability of cell-division and starting a new cell-cycle from state *g_i_*_1_ in the time interval (*t*, *t* + *dt*] is 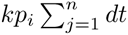

Theorem 1 of [55] gives the time derivative of the expected value of any function *φ*(*x*, *g_ij_*) as

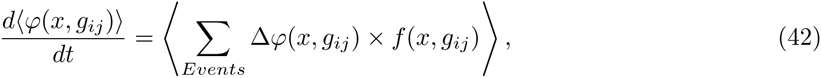

where Δ*φ*(*x*, *g_ij_*) is a change in *φ* when an event occurs. Based on this setup, mean dynamics of protein can be written by choosing *φ* to be *x*

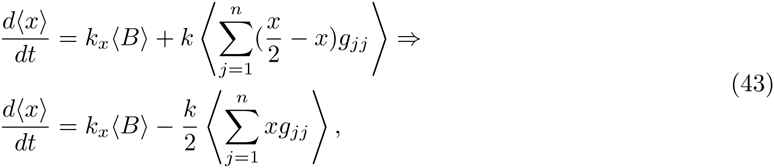

where we replaced conditional expected value of *x*_+_ by *x*/2 based on relation between statistical properties of *x*_+_ and *x* shown in equation (5).

Dynamics of 〈*x*〉 is not closed and depends to moments (*xg_ij_*), hence in order to have a closed set of equations we add new moments dynamics by selecting *φ* to be *xg_ij_* We do it in two steps: first we write the moment dynamics of 〈*xg*_11_〉

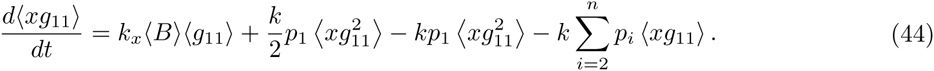

In the equation (9) of the main text it has been shown that

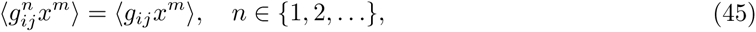

thus the term 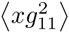 will simplify as

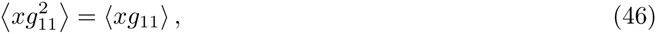

and the dynamics of 〈*xg*_11_〉 can be written as

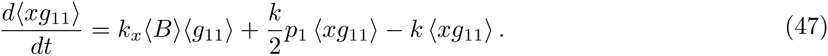

In the second step we write dynamics of the moments of the form 〈*xg_ij_*〉 other than 〈*xg*_11_〉

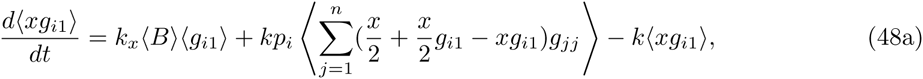

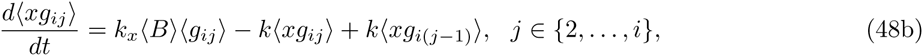

where dynamics of 〈*xg_i_*_1_〉 can be written as

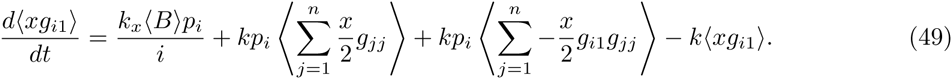

The equation (10) in the main text shows that

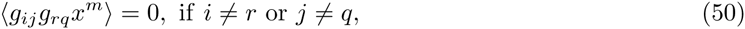

hence 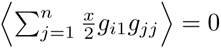, and equation (49) simplifies to

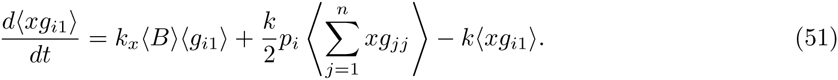

Further based on Figure 3 in the main text the probability of selecting a branch of *i* exponentials is *p_i_*, and because all the transitions happen with a constant rate *k*, hence mean of each of these *i* states is

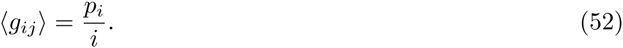

Thus equations (47), (48b), and (51) can be compactly written as shown in equation (11).

### B Moment dynamics of hybrid model introduced in Figure 2B

Stochastic hybrid system introduced in Figure 2B coupled with phase-type distribution contains the following stochastic events

| Event | Reset | Propensity |
| --- | --- | --- |
| Phase-type Evolution | $g_{ij}(t) \mapsto g_{ij}(t) - 1,$<br>$g_{i(j+1)}(t) \mapsto g_{i(j+1)}(t) + 1$ | $kg_{ij},$<br>$i \in \{2, \dots, n\},$<br>$j \in \{1, \dots, i-1\}$ |
| Cell-division | $x(t_s) \mapsto x(t_s)/2,$<br>$g_{jj}(t_s) \mapsto 0,$<br>$g_{i1}(t_s) \mapsto g_{i1}(t_s) + 1$ | $kp_i \sum_{j=1}^n g_{jj},$<br>$i \in \{1, \dots, n\}$ |

and deterministic protein production dynamics

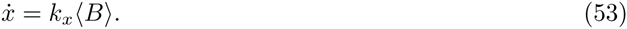

Time derivative of the expected value of any function *φ*(*x*, *g_ij_*) for this hybrid system can be written as [55]

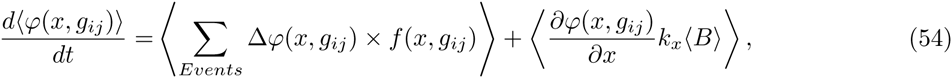

where the first term in the right-hand side is contributed from stochastic events and the second term is contributed from deterministic protein production dynamics. Based on this equation, the mean dynamics of the protein is calculated by choosing *φ* to be *x*

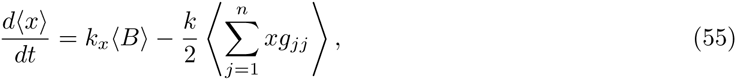

which is the same as equation (43). In addition to mean, dynamics of 〈*xg_ij_*〉 are also equal to their equation in the previous section.

The second order moment dynamics of protein can be expressed by choosing *φ* to be *x*^2^

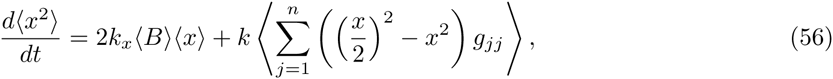

which can be simplified as

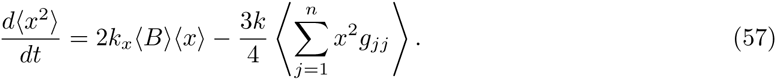

In order to have a closed set of equations we select *φ* to be of the form *x*^2^*g_ij_*. At the first step we write moment dynamics of 〈*x*^2^*g*_11_〉

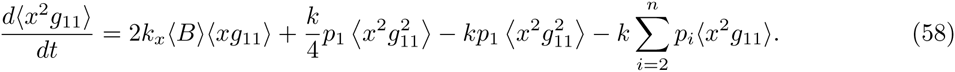

Based on equation (9) of the main text, the term 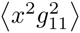 simplifies as

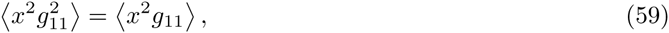

hence dynamics of 〈*x*^2^*g*_11_〉 will be

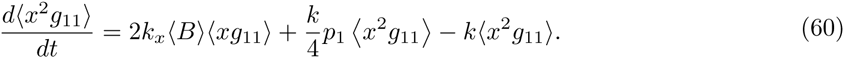

In the second step, we write dynamics of moments 〈*x*^2^*g_ij_*〉 when *g_ij_* ≠ *g*_11_

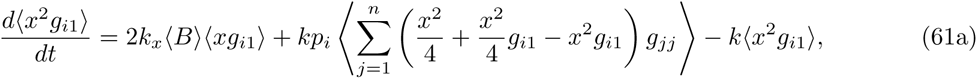

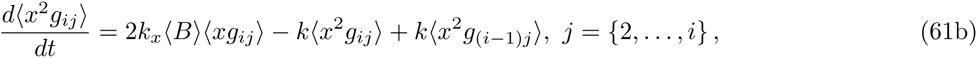

where dynamics of 〈*x*^2^*g_i_*_1_〉 can be shown to follow

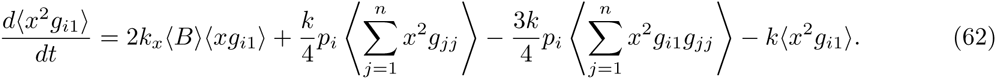

Based on equation (10) in the main text 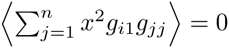, thus equation (62) simplifies to

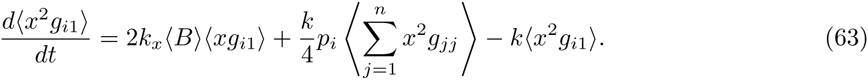

Equations (60), (61b), and (63) can be compactly written as equation (16) in the main text.

### C Moment dynamics of hybrid model introduced in Figure 2C

Stochastic hybrid system introduced in Figure 2C coupled with phase-type distribution contains the following stochastic events

| Event | Reset | Propensity |
| --- | --- | --- |
| Phase-type Evolution | $g_{ij}(t) \mapsto g_{ij}(t) - 1,$<br>$g_{i(j+1)}(t) \mapsto g_{i(j+1)}(t) + 1$ | $kg_{ij},$<br>$i \in \{2, \dots, n\},$<br>$j \in \{1, \dots, i-1\}$ |
| Cell-division | $x(t_s) \mapsto x_+(t_s),$<br>$g_{jj}(t_s) \mapsto 0,$<br>$g_{i1}(t_s) \mapsto g_{i1}(t_s) + 1$ | $kp_i \sum_{j=1}^n g_{jj},$<br>$i \in \{1, \dots, n\}$ |

and deterministic protein production dynamics

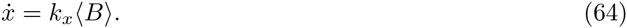

Note that in this model *x*(*t*) is a continuous random variable, thus we also use a continuous distribution to describe *x*_+_(*t_s_*), however statistical properties of *x*_+_(*t_s_*) is still given by (5). For this model we still can use equation (54) to derive moment dynamics; equations describing time evolution of mean and 〈*xg_ij_*〉 are the same as previous models, thus mean of protein for this model is equal to its value in Appendix A. The second order moment dynamics of protein can be written by choosing *φ* to be *x*^2^ in equation (54)

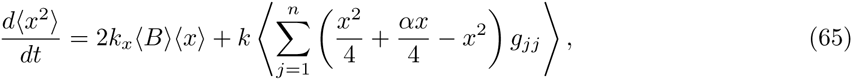

where conditional expected value of 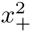 is substituted based on equation (5). Dynamics of 〈*x*^2^〉 can be simplified as

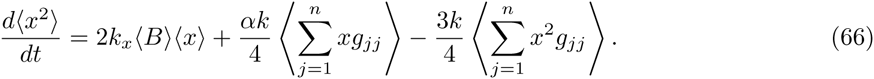

The same as before we add dynamics of the form 〈*x*^2^*g_ij_*〉 to have a closed set of dynamics. First we add dynamics of 〈*x*^2^*g*_11_〉

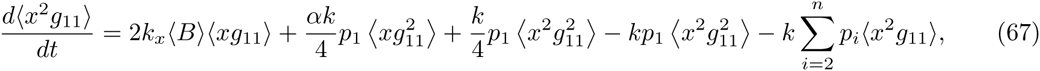

Based on equation (9) of the main text dynamics of 〈*x*^2^*g*_11_〉 simplifies to

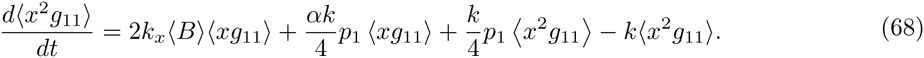

Now we express dynamics of moments 〈*x*^2^*g_ij_*〉 for *g_ij_* ≠ *g*_11_

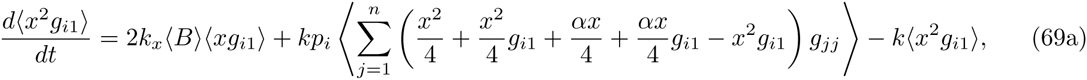

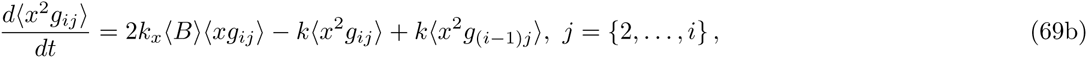

where dynamics of 〈*x*^2^*g_i_*_1_〉 can be shown as

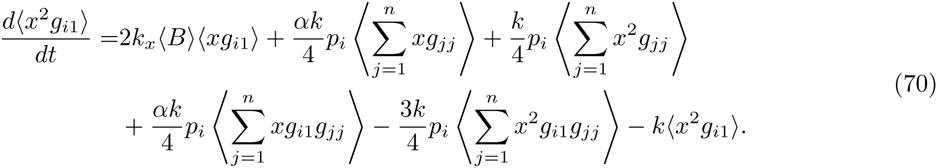

Based on equation (10) in the main text 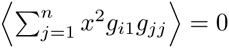, and 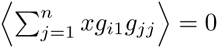, hence equation (70) simplifies to

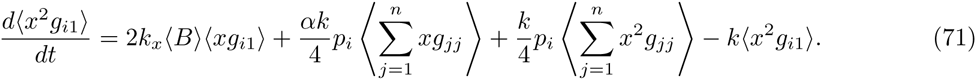

Equations (66), (68), (69b), and (71) can be compactly written as equation (23) in the main text.

### D Second and third-order moment dynamics of the full model

Based on model introduced in Appendix A, second order moment dynamics of protein is expressed by choosing *φ* to be *x*^2^ in equation (42),

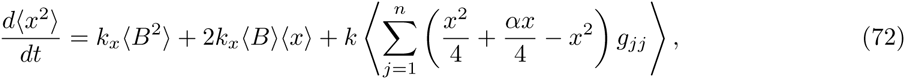

where conditional expected value of 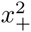 is substituted based on equation (5). Dynamics of 〈*x*^2^〉 can be simplified as

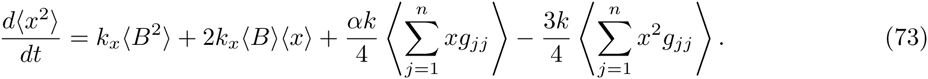

The same as before we add dynamics of the form 〈*x*^2^*g_ij_*〉 to have a closed set of moments. First we write
dynamics of 〈*x*^2^*g*_11_〉

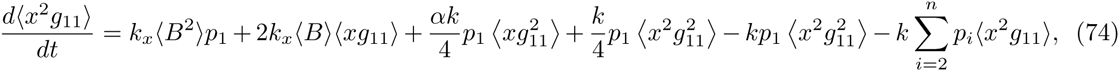

Based on equation (9) of the main text dynamics of 〈*x*^2^*g*_11_〉 simplifies to

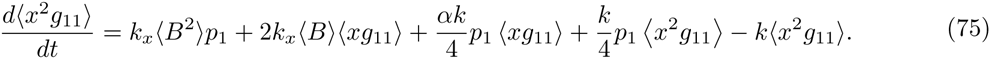

Next, dynamics of moments 〈*x*^2^*g_ij_*〉 when *g_ij_* ≠ *g*_11_ can be written as

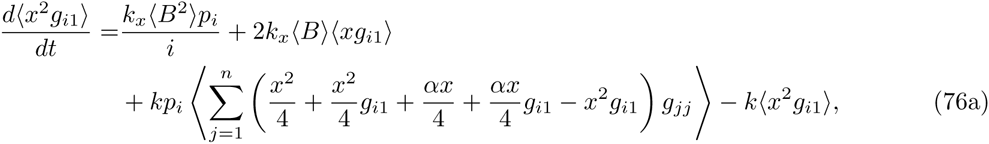

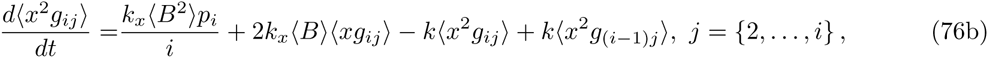

where dynamics of 〈*x*^2^*g*_i1_〉 can be shown as

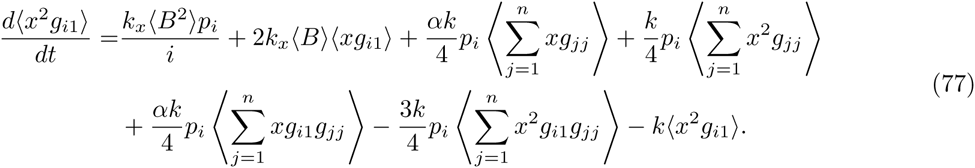

Based on equation (10) in the main text 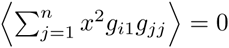 and 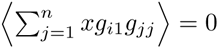, hence equation (77) simplifies to

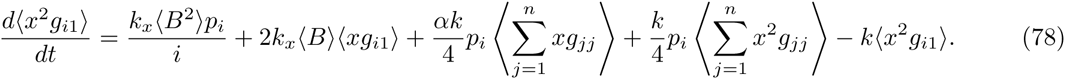

Equations (73), (75), (76b), and (78) can be compactly written as equation (26) in the main text.

### E Contribution of different sources of stochasticity in protein by taking into account gene-duplication

We study the contribution of different sources of stochasticity by using models introduced in Figure S1. The cell-cycle time consists of two time intervals: the time interval before gene-duplication and the time after gene-duplication. These time intervals are modeled by using two independent phase-type distributions as shown in Figure S2. Based on phase-type characteristics mean of the states of the first phase-type 〈*s_ij_*〉 and the second phase-type (*g_ij_*) are

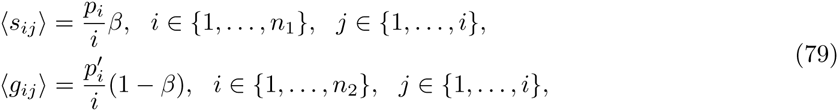

where *β* is defined as

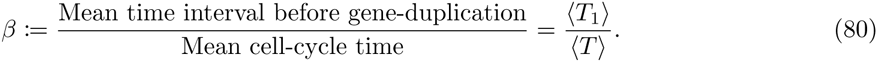

We start our analysis by deriving mean level of protein in the next section.

#### E.1 Mean of protein count level in the presence of gene-duplication

After gene-duplication the amount of genes expressing a specific protein doubles. Thus the rate of protein production increases by a factor of two as shown in Figure S1A. This model coupled with phase-type distributions contains the following stochastic events

Note that in the protein production event, before gene-duplication all the states *g_ij_* are zero thus propensity function will be 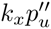. After gene-duplication and before division, one of the states *g_ij_* is one hence propensity function will be 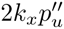 In time of gene-duplication, states of the first phase-type will reset to zero and state *g_i_*_1_ of the second distribution will be selected with probability 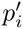; hence propensity function of gene-duplication event is 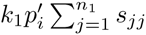. At the end of cell-cycle, states of the second phase-type will reset to zero and a new cell-cycle which is sum of *i* exponentials will be selected with probability *p_i_*; thus propensity function of cell-division event is 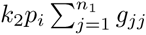.

**Figure S1:**
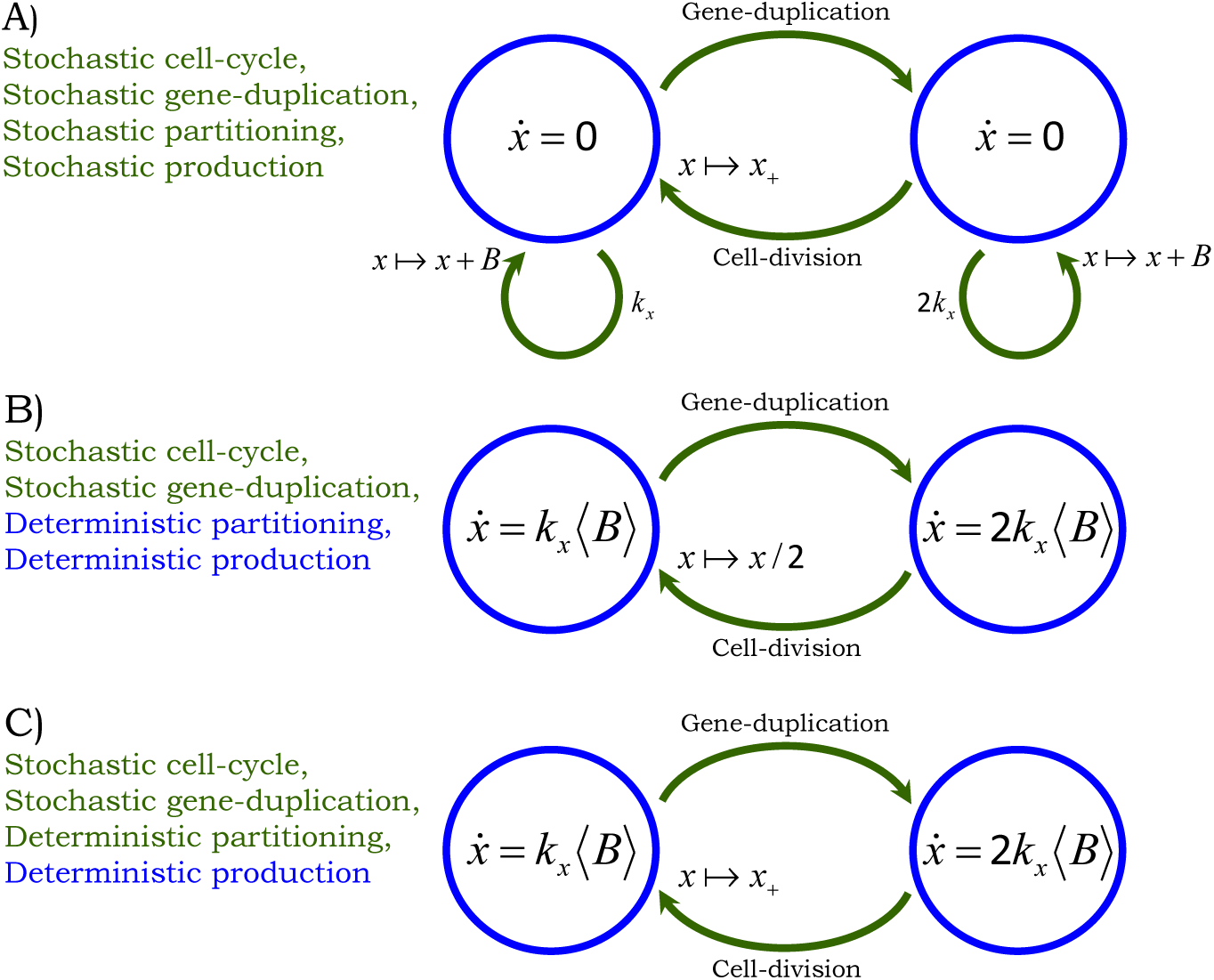
Stochastic hybrid models for quantifying different sources of noise. Gene-duplication and cell-division times are random events. **A)** Protein production happens in random bursts with burst frequency *k_x_*. After gene-duplication event burst frequency doubles (2*k_x_*). In the time of division proteins will be distributed between mother and daughter cells randomly, and the protein burst frequency will be *k_x_* again. **B)** Protein production is considered in a deterministic fashion, and after gene-duplication dynamics of protein production is multiplied by a factor of two, i.e., *ẋ* = 2*k_x_*〈*B*〉. In the division event proteins are distributed between mother and daughter cells equally. Thus the only stochastic events are duplication and division events. **C)** Protein is produced in a deterministic fashion, but in time of division protein levels in daughter and mother cells are random. Thus duplication, division, and partitioning are random events.

Theorem 1 of [55] gives the time derivative of the expected value of any function *φ*(*x*, s*_ij_*, *g_ij_*) as

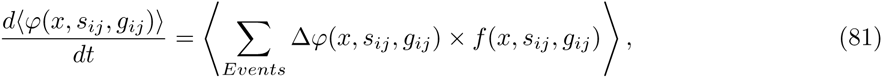

where Δ*φ*(*x*, s*_ij_*, *g_ij_*) is a change in *φ* when an event occurs. The first-order moment dynamic of this model can be expressed by selecting *φ* to be *x* in equation (81)

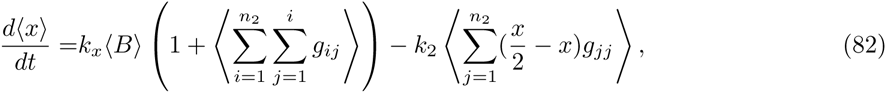

where conditional expected value of *x*_+_ is replaced from equation (5); by using equation (79) mean

**Figure S2:**
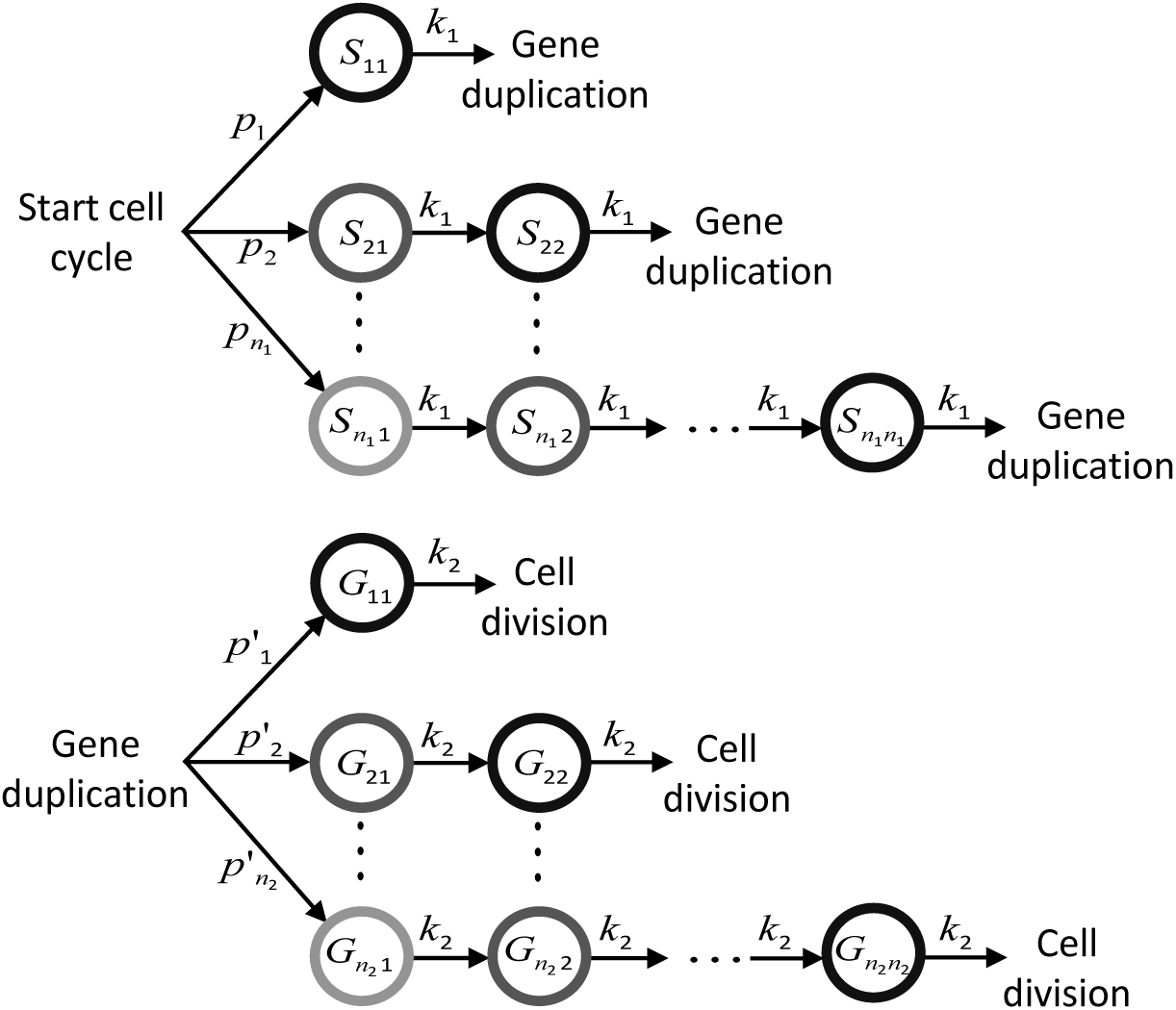
Cell-cycle time consists of two time intervals: at the end of the first interval gene duplicates, and at the end of the second one cell divides. Two independent phase-type distributions are used to model cell-cycle time in the presence of genome duplication. The states of the first distribution are denoted by *S_ij_*, *i* = {1,…, *n*_1_}, *j* = {1,…, *i*}; transition between these states happens at a constant rate *k*_1_. The states of the second distribution are shown by *G_ij_*, *i* = {1,…, *n*_2_}, *j* = {1,…, *i*}, and transition between these states occurs at a rate *k*_2_.

dynamics can be simplified as

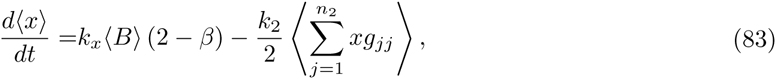

Mean dynamics is not closed thus we add dynamics of (*xs_ij_*), *i* = {1,…, *n*_1_}, *j* = {1,…, *i*} and 〈*xg_ij_*〉, *i* = {1,…, *n*_1_}, *j* = {1,…, *i*} to have a closed set of moment equations. These moment dynamics are simplified by using equations (5), (9), (10) and (79) as

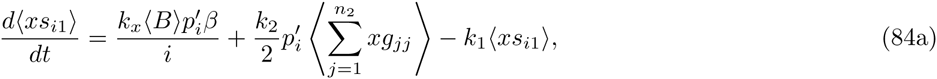

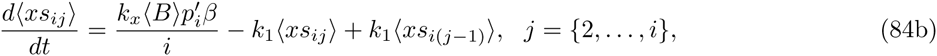

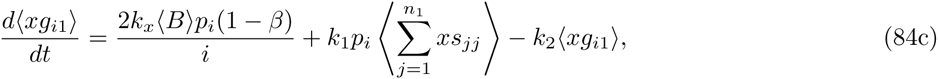

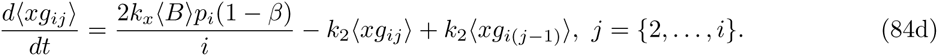

In order to find the mean of protein, first we need to find the moments 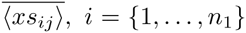, *i* = {1,…, *n*_1_}, *j* = {1, …, *i*} and 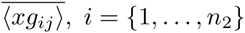, *i* = {1,…, *n*_2_}, *j* = {1, …, *i*}. For calculating these moments we should calculate the term 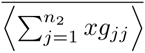; this term can be obtained by analyzing equation (83) in steady-state

| Event | Reset | Propensity |
| --- | --- | --- |
| Protein production | $x(t) \mapsto x(t) + u$ | $k_x p_u'' \left( 1 + \sum_{i=1}^{n_2} \sum_{j=1}^i g_{ij} \right)$ |
| First phase-type evolution | $s_{ij}(t) \mapsto s_{ij}(t) - 1,$<br>$s_{i(j+1)}(t) \mapsto s_{i(j+1)}(t) + 1$ | $k_1 s_{ij},$<br>$i \in \{2, \dots, n_1\},$<br>$j \in \{1, \dots, i-1\}$ |
| Gene-duplication | $s_{jj}(t) \mapsto 0,$<br>$g_{i1}(t) \mapsto g_{i1}(t) + 1$ | $k_1 p_i' \sum_{j=1}^{n_1} s_{jj},$<br>$i \in \{1, \dots, n_2\}$ |
| Second phase-type evolution | $g_{ij}(t) \mapsto g_{ij}(t) - 1,$<br>$g_{i(j+1)}(t) \mapsto g_{i(j+1)}(t) + 1$ | $k_2 g_{ij},$<br>$i \in \{2, \dots, n_2\},$<br>$j \in \{1, \dots, i-1\}$ |
| Cell-division | $x(t_s) \mapsto x_+(t_s),$<br>$g_{jj}(t_s) \mapsto 0,$<br>$s_{i1}(t_s) \mapsto s_{i1}(t_s) + 1$ | $k_2 p_i \sum_{j=1}^{n_2} g_{jj},$<br>$i \in \{1, \dots, n_1\}$ |

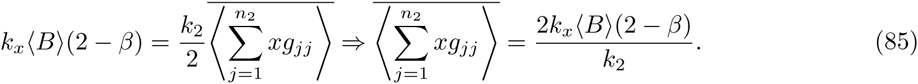

By having this term, we calculate 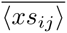 by recursion process: we start by calculating 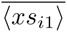 by substituting equation (85) in equation (84a). In the next step we use the definition we derived for 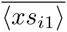 to calculate 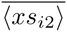 from equation (84b). We continue this process until we derive all the moments

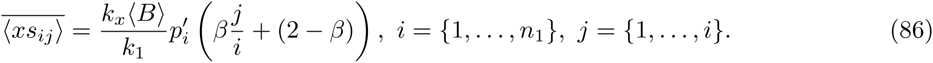

Now we need to calculate the moments 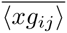, *i* = {1,…, *n*_2_}, *j* = {1,…, *i*}, thus we need the expression of the term 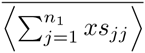; from equation (86) we have the following

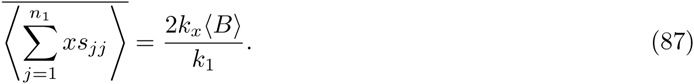

Substituting this term in equations (84c) and (84d) result in

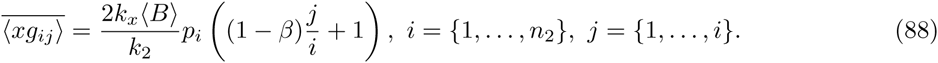

Note that

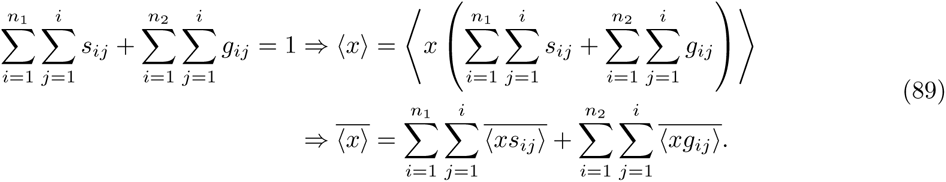

Thus by adding all the term calculated here and using equation (7) mean of protein can be calculated as

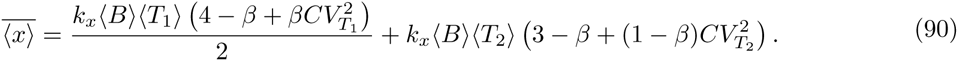

#### E.2 Noise in protein count level contributed from cell-cycle time

In order to calculate the noise contributed from cell-cycle time variation, the model introduced in Figure S1B coupled with phase-type distributions is used. This model contains following stochastic events

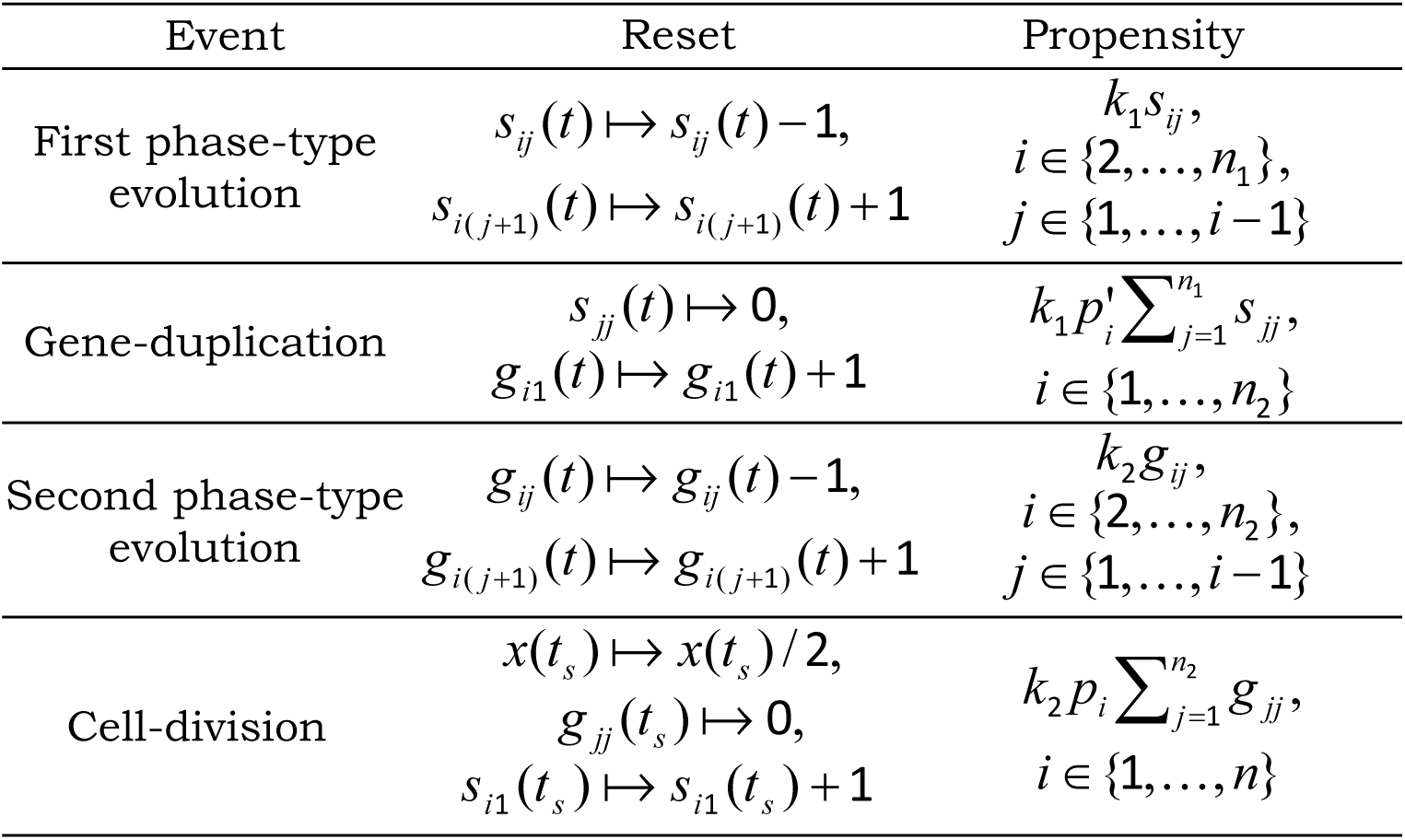

and deterministic protein production

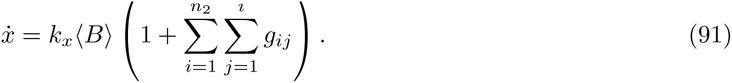

Theorem 1 of [55] gives the time derivative of the expected value of any function *φ*(*x*, *s_ij_*, *g_ij_*) as(92)

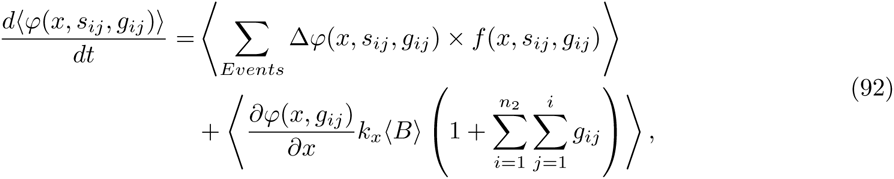

where the first term in the right hand side is contributed from stochastic events, and the second term is contributed from deterministic protein production. In this model, dynamics of 〈*x*〉, 〈*xs_ij_*〉 and 〈*xg_ij_*〉 are the same as equations (83) and (E.6), thus mean of protein, 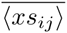, and 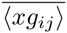 will be equal to their value in previous section. Further, the second-order moment dynamics of protein can be added by selecting *φ* to be *x*^2^ in equation (92)

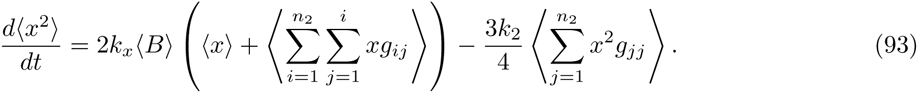

This equation is not closed thus we add dynamics of 〈*x*^2^*s_ij_*〉, *i* = {1,…,*n*_1_}, *j* = {1,…,*i*} and 〈*x*^2^*s_ij_*〉, *i* = {1,…,*n*_2_}, *j* = {1,…,*i*} to have a closed set of equations

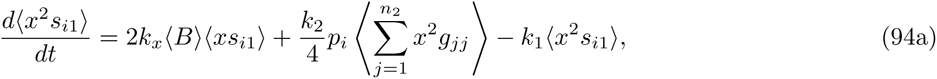

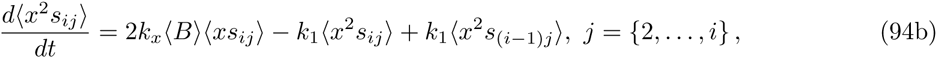

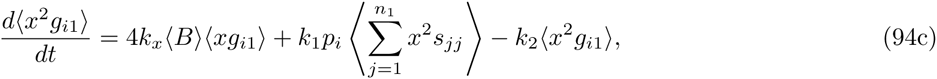

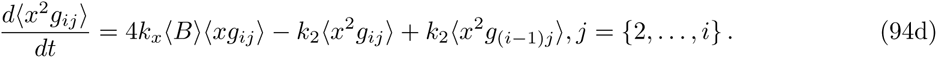

In order to calculate noise we need to express 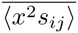, and 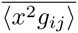, which requires calculating the term 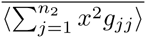; this term can be derived by analyzing equation (93) in steady-state

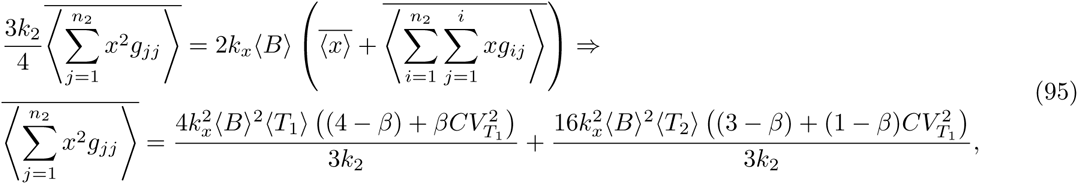

where in deriving this term we used equation (90) and we summed all the terms in equation (88). By having this term, we calculate 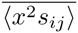 by recursion process: we derive 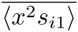 by substituting equation (95) in equation (94a). In the next step we use the definition of 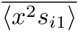 to calculate 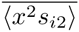 from equation (94b). We continue this process until we derive all the moments

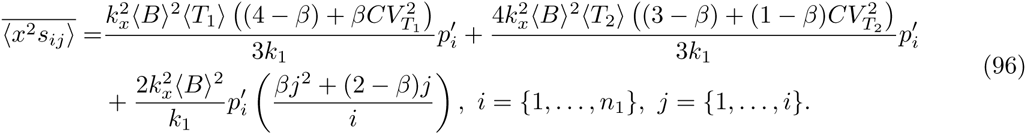

Expressing 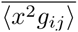 requires calculation of the term 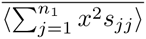 which can be obtained from equation

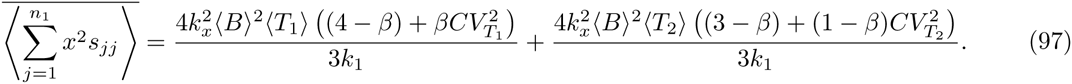

Thus 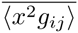 can be obtained with a recursion process from equations (94c) and (94d)

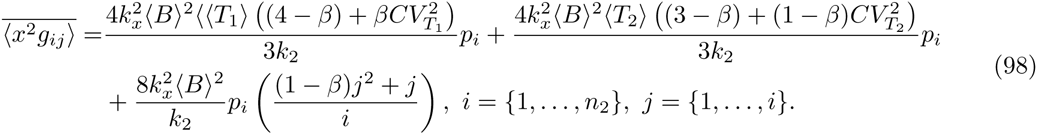

Note that 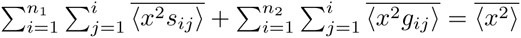 thus the second order moment of protein can be derived by adding all the terms in equations (96) and (98). 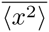 can be simplified by using equations (7) and (18b) in the main article as

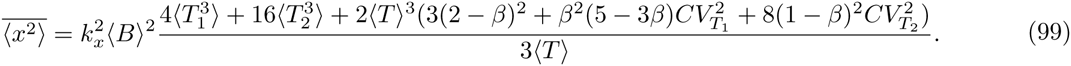

Finally, using the definition of *CV*^2^ results in noise of protein raised from cell-cycle time variations

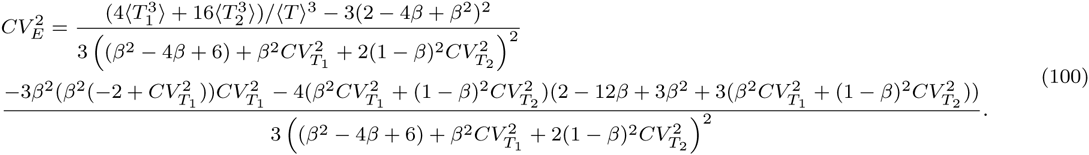

#### E.3 Noise in protein count level contributed from random partitioning

In order to take into account noise caused by random partitioning of proteins between two daughter cells, we use the model shown in Figure S1C coupled with phase-type distributions. This model contains the following stochastic events

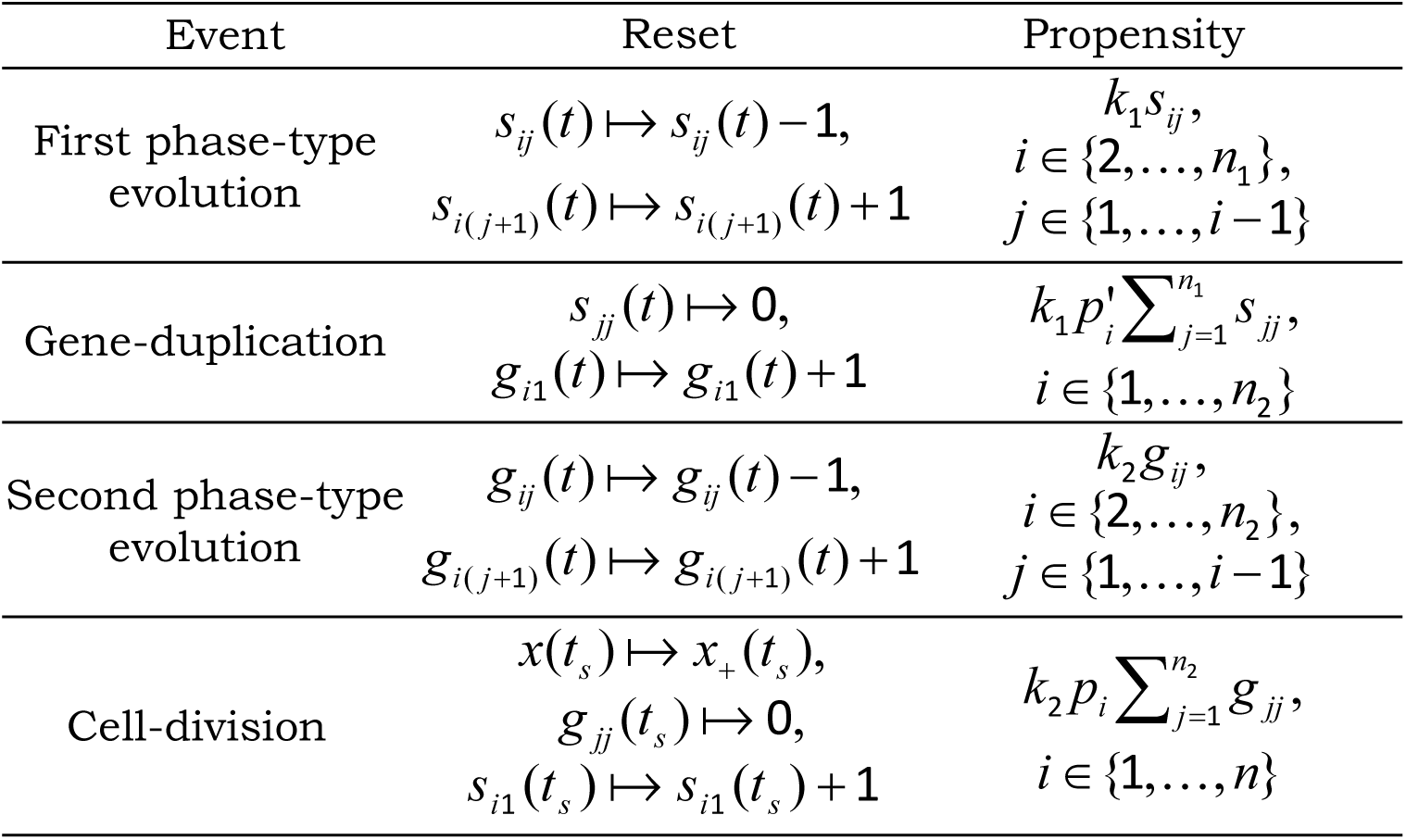

| Event | Reset | Propensity |
| --- | --- | --- |
| First phase-type evolution | $s_{ij}(t) \mapsto s_{ij}(t) - 1,$<br>$s_{i(j+1)}(t) \mapsto s_{i(j+1)}(t) + 1$ | $k_1 s_{ij},$<br>$i \in \{2, \dots, n_1\},$<br>$j \in \{1, \dots, i-1\}$ |
| Gene-duplication | $s_{jj}(t) \mapsto 0,$<br>$g_{i1}(t) \mapsto g_{i1}(t) + 1$ | $k_1 p'_i \sum_{j=1}^{n_1} s_{jj},$<br>$i \in \{1, \dots, n_2\}$ |
| Second phase-type evolution | $g_{ij}(t) \mapsto g_{ij}(t) - 1,$<br>$g_{i(j+1)}(t) \mapsto g_{i(j+1)}(t) + 1$ | $k_2 g_{ij},$<br>$i \in \{2, \dots, n_2\},$<br>$j \in \{1, \dots, i-1\}$ |
| Cell-division | $x(t_s) \mapsto x_+(t_s),$<br>$g_{jj}(t_s) \mapsto 0,$<br>$s_{i1}(t_s) \mapsto s_{i1}(t_s) + 1$ | $k_2 p_i \sum_{j=1}^{n_2} g_{jj},$<br>$i \in \{1, \dots, n\}$ |

and deterministic protein production

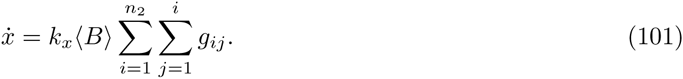

Note that here *x* is a continuous random variable, hence *x*_+_ is also obtained from a continious distribution. Connection between statistical statistical moments of *x* and *x*_+_ is given by (5).

For this model, 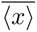, 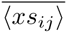, and 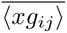 are equal to their value in Section E.1 and Section E.2. However, dynamics of 〈*x*^2^〉 and (*x*^2^*s_i_*_1_) are different

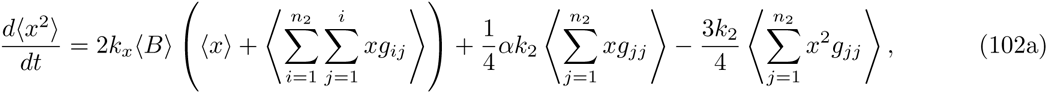

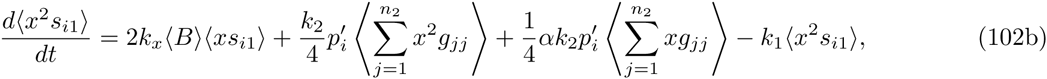

note that dynamics of 〈*x*^2^*s_ij_*〉, 〈*x*^2^*g*_i1_〉, and 〈*x*^2^*g_ij_*〉 are identical to equations (94b), (94c), and (94d). Similar to previous section, we start by deriving the term 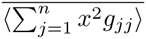. Analyzing equation (102a) in steady-state gives this term as

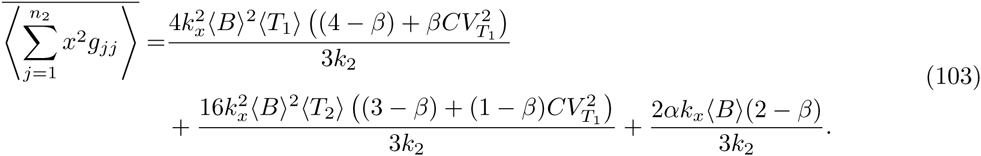

Substituting equation (103) in equations (102b) and (94b) results in

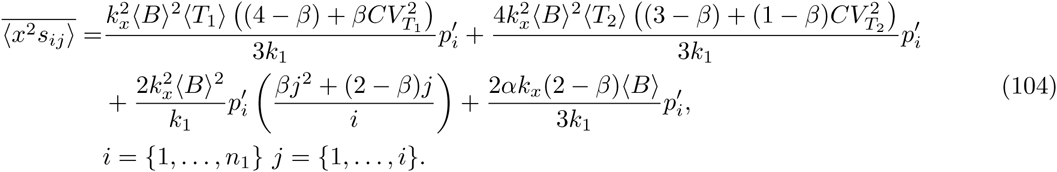

In the next step we derive moments 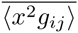; we start by calculating 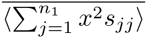 from (104)

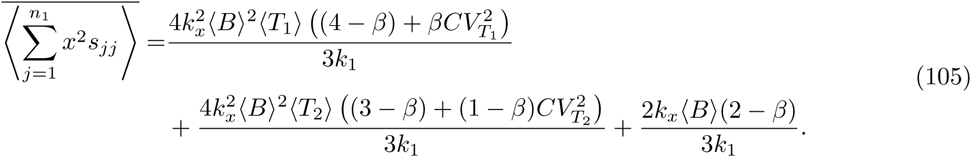

By having this term, the moments 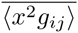 are derived by solving equations (94c) and (94d) in steady-state

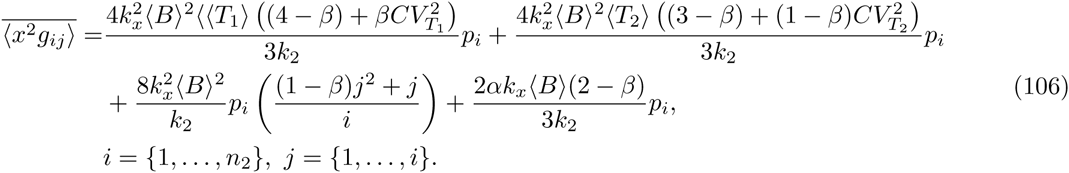

Note that

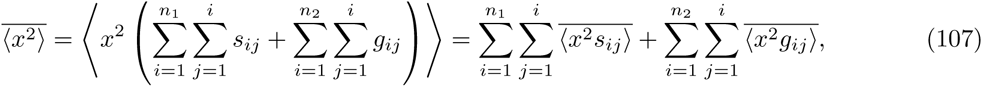

hence the second-order moment is

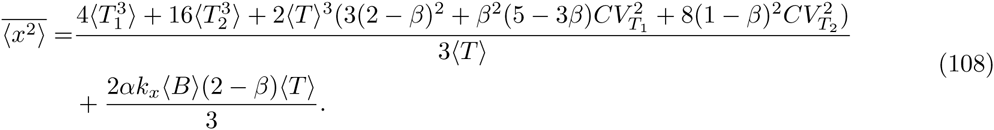

Coefficient of variation squared gives noise raised from partitioning and cell-cycle variations, which subtracting equation (100) from results gives partitioning noise as

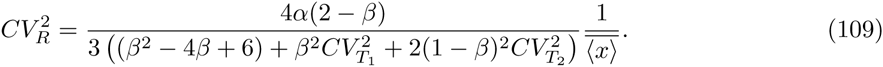

#### E.4 Noise in protein count level contributed from stochastic production

In order to calculate the noise caused by stochastic birth of protein, we use the model introduced in Section C.1. For this model, moments dynamics of 〈*x*^2^〉, 〈*x*^2^*s_ij_*〉, and 〈*x*^2^*g_ij_*〉 can be written as

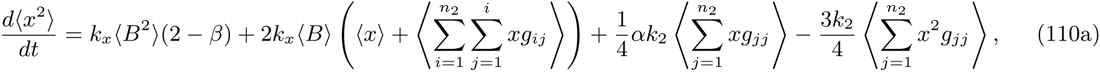

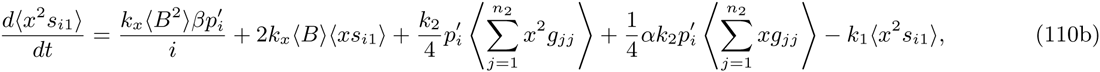

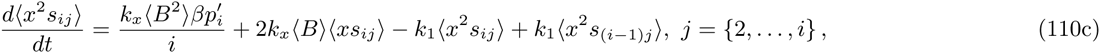

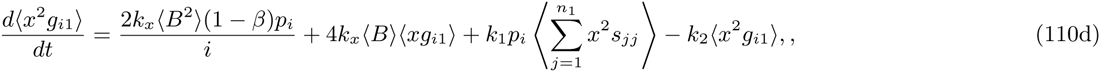

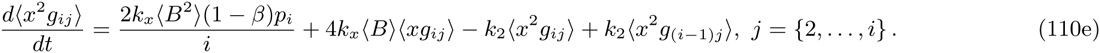

The same as before we start by expressing the term 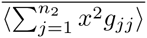, this term is calculated by analyzing equation (110a) in steady-state

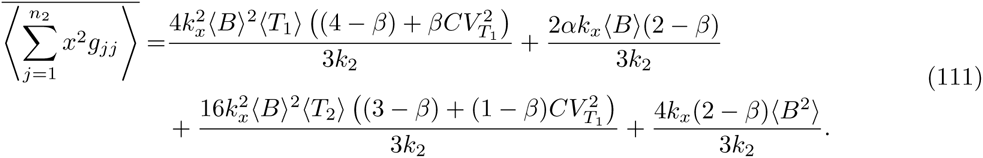

Substituting this term in equations (110b) and (110c) results in

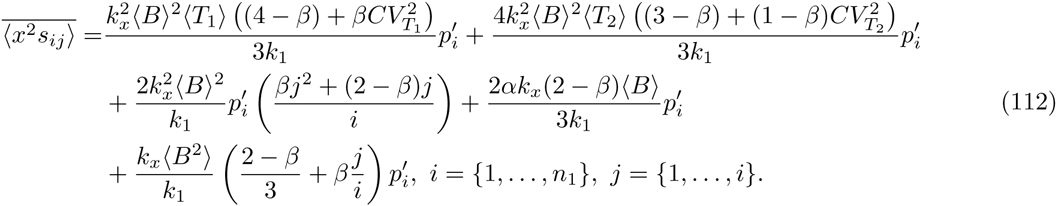

Similar to previous section, solving equations (110d) and (110e) gives the 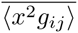

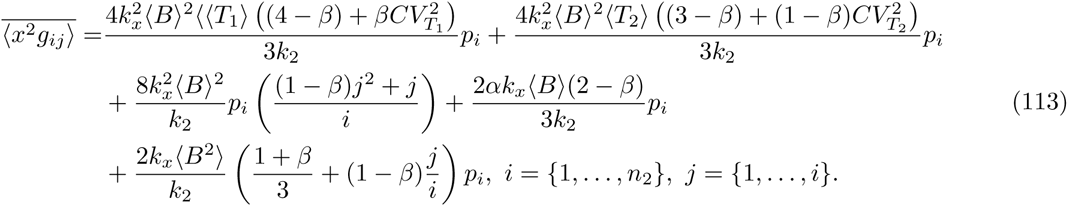

Finally summing all the moments 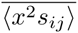, and 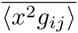 results in 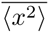 as

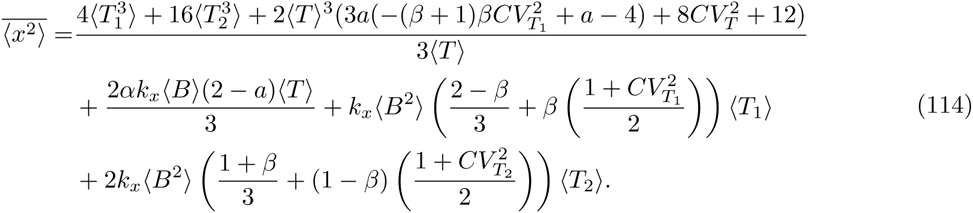

Steady-state analysis gives the noise from stochastic birth, random partitioning, and cell-cycle time variations. Subtracting noise of cell-cycle time and partitioning in equations (100) and (109) results in noise caused by stochastic production of protein

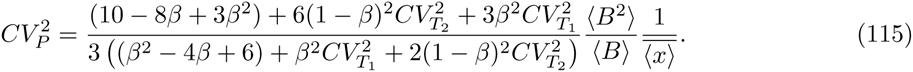

#### E.5 Effect of gene-duplication time in intrinsic noise

We investigate how the noise contributions from random partitioning and stochastic expression (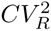 and 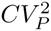 terms in equation (34) of the main text) change as *β* is varied between 0 and 1. Results show that 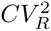 and 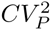 follow the same qualitative shapes as reported in Fig. 6. There exists a *β*\*

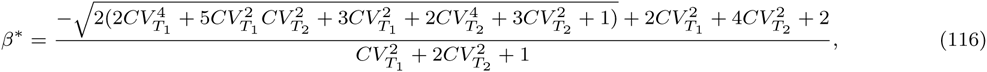

such that 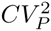 is minimized and 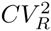 is maximized when *β* = *β*\*. Note that when 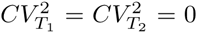, 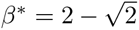 as reported in the main text. The minimum value of 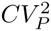 and the maximum value of 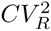 are given by

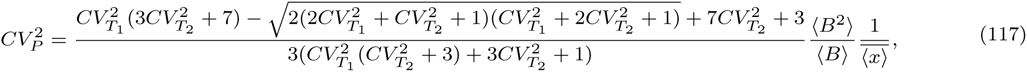

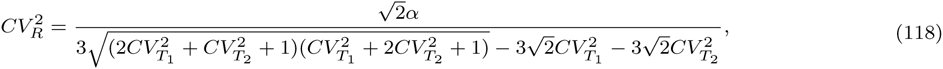

respectively. Plots of *β*\* and optimal value of 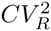 and 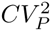 as a function of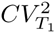are shown in Fig. S4. Note that if noise in *T*_1_ is high and *T*_2_ is deterministic then *β*\* shifts towards zero. Similarly, if noise in *T*_2_ is high and *T*_1_ is deterministic then *β*\* shifts towards one.

### F Noise level in unstable protein

Consider an unstable protein with sufficiently high degradation rate *γ_x_* such that the protein level reaches steady-state instantaneously compared to the cell-cycle time (Fig. S4). Let *τ* denote the time from the last division event, then

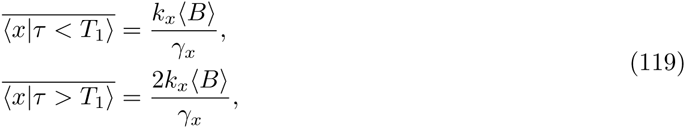

where *T*_1_ is the time in which duplication happens. The mean level of an unstable protein can be calculated as

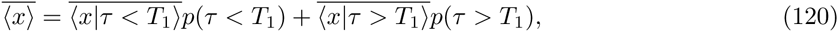

**Figure S3:**
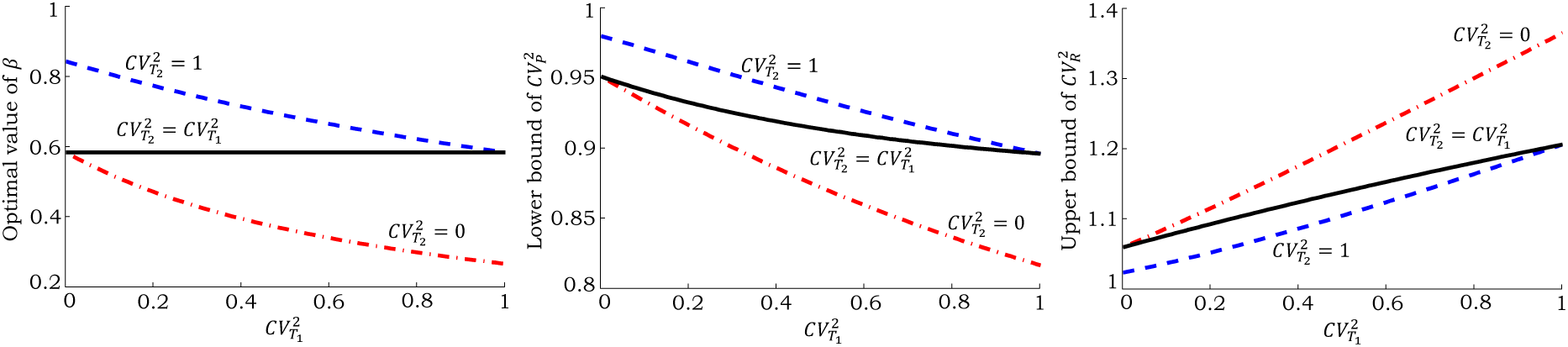
Effect of gene-duplication on intrinsic noise level. *Left*: Value of *β* where 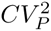 is minimized and 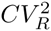 is maximized as a function of 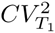. When 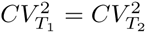, noise levels always reach their extrema at 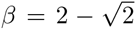. *Middle & Right*: Extremum values of 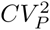 and 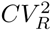 as a functions of 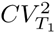. Noise levels are normalized by their values at *β* = 0.

where *p*(*τ* < *T*_1_) and *p*(*τ* > *T*_1_) denote the probability of being in the time interval before and after gene-duplication. Using

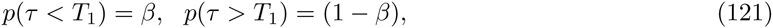

we obtain

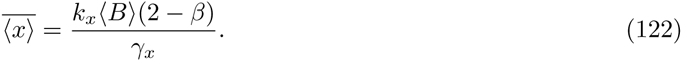

To compute the extrinsic noise component we consider deterministic protein production and decay. The second-order moment of *x*(*t*) is given by

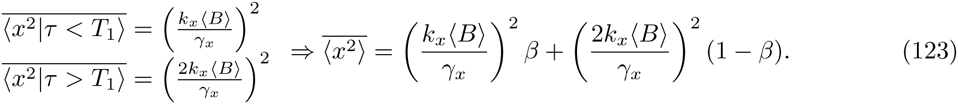

By using definition of *CV*^2^, extrinsic noise is

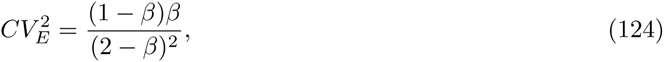

which is zero at *β* = 0,1 and reaches its maximum at *β* = 2/3 (Fig. S4).

Next we compute the intrinsic noise component. If the protein decay is sufficiently high, the noise contribution from partitioning errors will be negligible because any errors will be instantaneously corrected due to rapid protein turnover. Noise raised from stochastic gene expression can be investigated by considering a model containing stochastic bursty production and stochastic degradation of proteins, where after gene-duplication the burst frequency doubles. Again assuming large enough 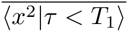 is equal to the steady-state second-order moment of a stochastic model with burst frequency *k_x_* (analyzed in [64])

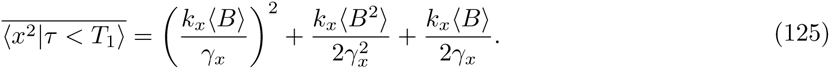

In comparison with equation (123), there are two extra terms at the right hand side of 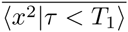. The first extra term is due to production of protein in random bursts and the second one is due to stochastic degradation of protein molecules. Further for the same reasons (large degradation rate and rapid equilibration of the distribution), 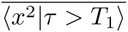 is equal to the second-order moment of a model containing stochastic bursty production of proteins with burst frequency 2*k_x_* which is

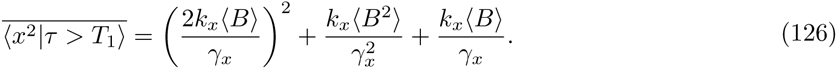

Thus the second order moment of an unstable protein can be written as

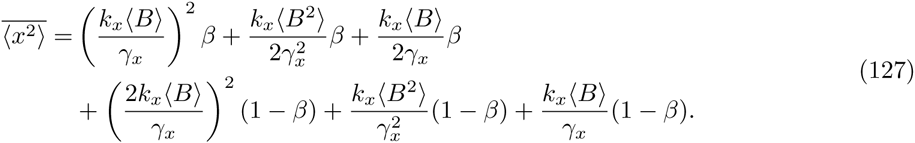

Using definition of *CV*^2^ and subtracting extrinsic noise we obtain the following noise contribution from stochastic expression and decay

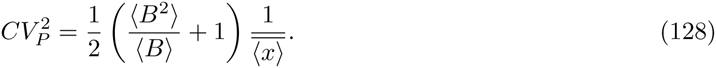

**Figure S4:**
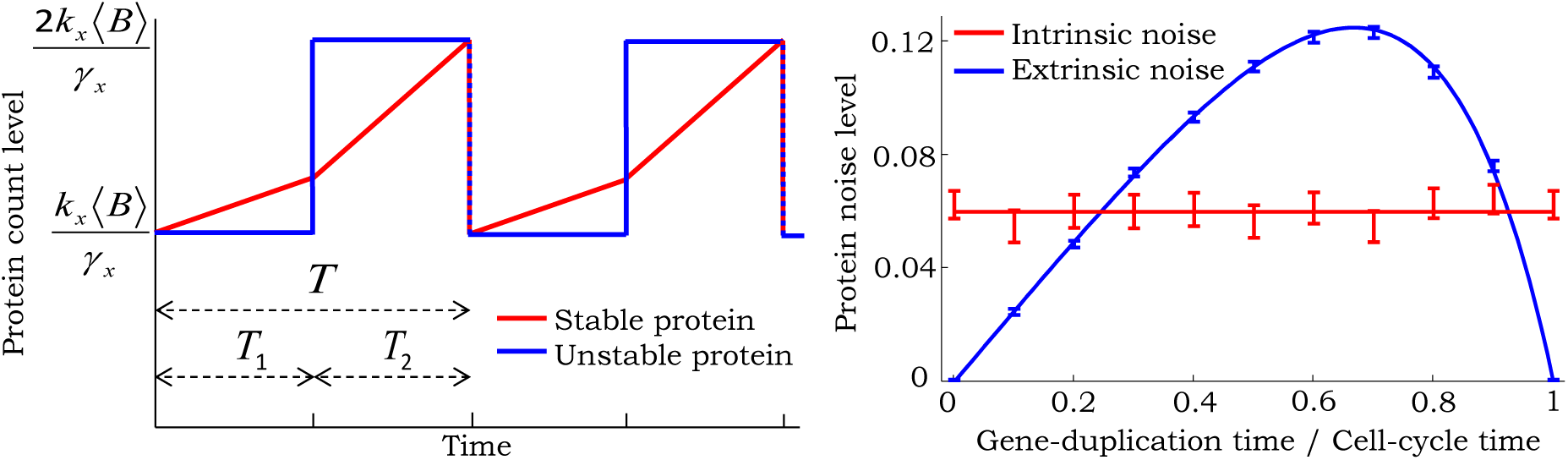
Contribution of gene duplication to noise levels of an unstable protein. *left*: For a stable protein, copy numbers accumulate in a bilinear fashion. In contrast, an unstable protein reaches equilibrium rapidly and its level changes in steps. *Right*: Extrinsic and intrinsic noise predicted for an unstable protein as a function of *β*. Solid lines are predictions from (124) and (128), which agree with estimates from 20, 000 Monte Carlo simulations. Parameters taken as *γ_x_* = 10*hr*^−1^, and a geometric burst with 〈*B*〉 = 6. Burst frequency is changed to have a constant mean protein level of 100 molecules for different values of *β*. 95% confidence intervals are calculated via bootstrapping.

## Acknowledgements

AS is supported by the National Science Foundation Grant DMS-1312926.

